# Phenotypic dissection of epithelial lineages and therapeutic manipulation of differentiation programs in human Adenoid Cystic Carcinomas (ACCs)

**DOI:** 10.1101/2022.01.19.476843

**Authors:** Sara Viragova, Luis Aparicio, Junfei Zhao, Luis E. Valencia Salazar, Alexandra Schurer, Anika Dhuri, Debashis Sahoo, Christopher A. Moskaluk, Raul Rabadan, Piero Dalerba

## Abstract

*Adenoid Cystic Carcinoma* (ACC) is a rare and aggressive form of salivary gland cancer, characterized by the co-existence within tumor tissues of two distinct populations of malignant cells, phenotypically similar to the myoepithelial and ductal lineages of normal salivary glands. Using a novel computational approach for single-cell RNA-seq analysis, we identified two cell-surface markers (CD49f, KIT) that enable the differential purification of myoepithelial-like (CD49f^high^/KIT^neg^) and ductal-like (CD49f^low^/KIT^+^) cells from ACC *patient derived xenografts* (PDX). Using prospective xeno-transplantation experiments, we demonstrate that myoepithelial-like cells act as progenitors of ductal-like cells. Using *three-dimensional* (3D) organoid cultures, we demonstrate that agonists of *retinoic acid* (RA) signaling promote differentiation of myoepithelial-like cells into ductal-like cells, while inhibitors of RA signaling selectively kill ductal-like cells. Finally, we demonstrate that BMS493, an inverse agonist of RA signaling, can be successfully leveraged for the *in vivo* treatment of human ACCs.

## INTRODUCTION

*Adenoid cystic carcinomas* (ACCs) are malignant adenocarcinomas which arise in secretory glands throughout the body, most commonly in the *salivary glands* (SGs) of the head and neck^1^. ACCs usually display an indolent growth, but their slow proliferation kinetics belie their biologically aggressive and clinically relentless nature, characterized by high rates of local invasion, peri-neural infiltration, and high frequency of distant metastases, often 10-15 years after initial diagnosis^2^.

Current treatment options for ACCs are limited to surgery and radiotherapy, which are often destructive and, in approximately 60% of cases, unable to prevent metastatic relapse and patient death. Due to their slow proliferation kinetics and low rates of somatic mutation (i.e., low mutational burdens), ACCs are usually refractory to chemotherapy and immunotherapy regiments^3, 4^ and lack meaningful targets for the development of pharmacological agents with tumor-specificity (i.e., kinase inhibitors)^5, 6^.

From a histological point of view, ACCs are characterized by a defining feature: the concurrent presence of two phenotypically distinct populations of cancer cells, termed “ductal-like” and “myoepithelial-like” because of their similarity to the ductal and myoepithelial cell lineages of the normal SG epithelium, in terms of both morphology and expression of lineage-specific biomarkers^1, 7–9^. Although this feature has long been appreciated, the molecular mechanisms that underpin it remain poorly understood. It remains unclear, for example, whether the two cell types originate as a result of clonal events (i.e., the accumulation of divergent somatic mutations) or of multi-lineage differentiation processes akin to those that enable stem/progenitor cells to sustain the homeostatic turnover of adult tissues^10^, as in the case of colon cancer^11^. In the case of ACC, this important aspect of the tumor’s biology has remained difficult to investigate, due to the lack of experimental means that would enable the isolation of the two cell populations from primary tumors, and allow their comparison in terms of transcriptional identities, functional properties, developmental relationships, and differential sensitivity to anti-tumor therapies.

In this study, we utilize *single cell RNA sequencing* (scRNA-seq) in concert with a novel computational approach derived from *random matrix theory* (RMT)^12^, to identify cell surface markers that enable, for the first time, the differential purification by *fluorescence activated cell sorting* (FACS) of the two malignant cell populations found in *patient-derived xenograft* (PDX) lines established from human ACCs^13^. We then proceed to test their developmental relationships using prospective transplantation experiments, to identify a signaling pathway that controls the differentiation of one into the other, and to develop a new therapeutic approach for the *in vivo* treatment of human ACCs.

## RESULTS

### Identification of cell-surface markers for the differential purification of myoepithelial-like and ductal-like cell types from human ACCs

To identify cell-surface markers differentially expressed between the myoepithelial-like and ductal-like populations of human ACCs, we performed scRNA-seq on the PDX line ACCX22. We selected this model because it was established from a parotid gland ACC lesion, representing one of the most frequent sites of origin for ACCs^8^, and because it displayed two classical morphological features of human ACCs: a well-differentiated “cribriform” histology and a bi-phenotypic cell composition. Utilizing our previously established protocols for dissociation and single-cell transcriptional analyses of PDX tumors^11, 14^, we dissociated ACCX22 into a single-cell suspension, purified human epithelial (EpCAM^+^) tumor cells by FACS, and used scRNA-seq to capture their individual transcriptional profiles **(Extended Data Fig. 1)**. We then implemented a novel computational pipeline (*Randomly*)^12^ designed to leverage the statistical principles of RMT to remove gene-expression signals that follow a stochastic distribution and identify genes most responsible for biological signal. We clustered cells in different subgroups based on transcriptional patterns **(Extended Data Fig. 2)**, with the optimal clustering solution (i.e., the one displaying the highest silhouette score) consisting of three sub-populations **(Figure 1a)**. The two most abundant populations displayed mutually exclusive expression of known myoepithelial-like (e.g., *ACTA2*, *CNN1*, *TP63*) and ductal-like (*KRT7, KRT18, ELF5*) markers **(Extended Data Fig. 3a-b)**, while a third population appeared to represent a highly proliferating (*MKI67*^high^, *TOP2A*^high^*, CDK1*^high^, *PCNA*^high^) subset of the population expressing ductal-like markers **(Figure 1e, Extended Data Fig. 3c)**. Among the genes displaying the highest differential expression between the two main clusters, we identified cell-surface markers CD49f (*ITGA6*), and KIT/CD117 (*KIT*), which were preferentially detected in cells associated with myoepithelial-like and ductal-like markers, respectively **(Figure 1b-d)**. We then tested whether the expression levels of CD49f and KIT proteins could be leveraged to visualize the two subsets of malignant cells using FACS. Indeed, the combined use of the two markers allowed us to discriminate two clearly distinct cell populations (CD49f^high^/KIT^neg^ vs. CD49f^low^/KIT^+^), across 5 independent PDX lines representative of bi-phenotypic ACCs (ACCX5M1, ACCX6, ACCX14, ACCX22, SGTX6) **(Figure 1f)**. Analysis of the same tumors by *immunohistochemistry* (IHC) confirmed that KIT expression was restricted to cells with ductal-like morphology, and mutually exclusive to expression of TP63, a marker of myoepithelial-like cells, in agreement with previous reports^9^ **(Figure 1g)**.

**Figure 1:**
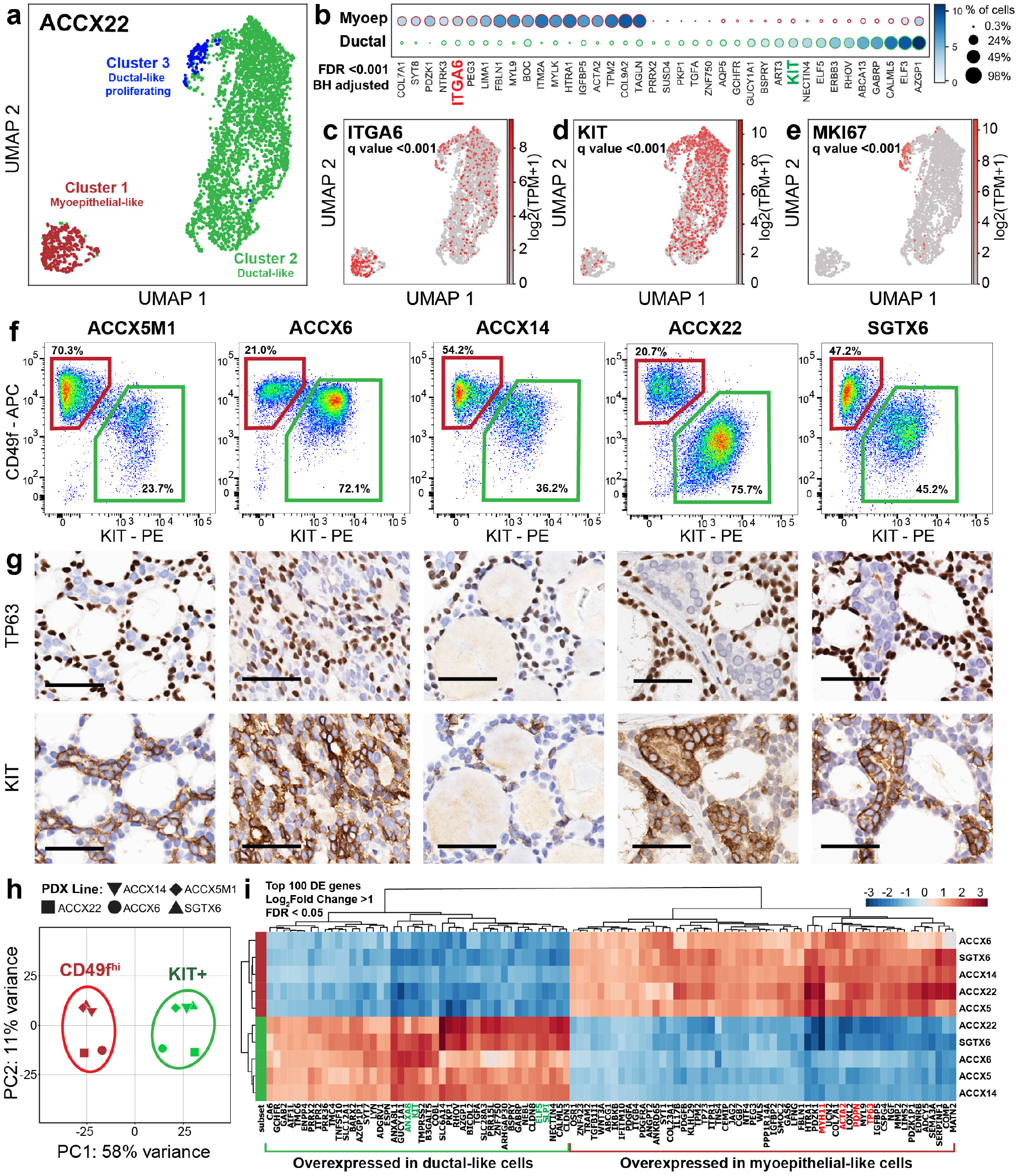
Identification of surface markers for the differential purification of myoepithelial-like and ductal-like populations from human ACCs. **(a)** Visualization of scRNA-seq data from PDX line ACCX22 using *Uniform Manifold Approximation and Projection* (UMAP). The UMAP scatter plot reveals three major cell clusters: 1) myoepithelial-like cells (red); 2) ductal-like cells (green); and 3) ductal-like cells undergoing mitosis (blue). **(b)** List of genes displaying the most robust difference in mean expression levels between myoepithelial-like cells (Cluster 1) and ductal-like cells (Cluster 2), after testing for statistical significance (Student’s t-test) and adjustment for multiple comparisons (FDR<0.001; Benjamini-Hochberg method). Among the top differentially expressed genes are two surface markers: ITGA6 (CD49f) and KIT (CD117). **(c-e)** UMAP plots displaying gene expression levels for ITGA6 (c), KIT (d) and the proliferation marker MKI67 (e). **(f)** Analysis by flow cytometry of five human PDX lines representative of bi-phenotypic ACCs, revealing the co-existence of two distinct human (mCd45^neg^/mH2-kd^neg^), epithelial (EpCAM+) cancer cell populations, defined by differential expression of surface markers CD49f and KIT. **(g)** Analysis by *immunohistochemistry* (IHC) of PDX tumor tissues, confirming the coexistence of two sub-populations of malignant cells, defined by mutually exclusive expression of TP63 (a myoepithelial marker) and KIT (a ductal marker). Scale bar = 50 µm. **(h)** *Principal component analysis* (PCA) of RNA-seq data obtained from autologous pairs of CD49f^high^/KIT^neg^ (red) and CD49f^low^/KIT^+^ (green) cells, purified in parallel from five PDX lines, revealing tight clustering of the two cell types along the first principal component (PC1). **(i)** Heatmap visualization of the top 100 differentially expressed genes between CD49f^high^/KIT^neg^ and CD49f^low^/KIT^+^ cells, after mean-centering of gene-expression levels and hierarchical clustering of both genes and samples. Differentially expressed genes were defined as genes with a more than two-fold difference in mean expression levels between the two populations (log2 fold change = 1) that was considered to be statistically robust after ranking based on the Wald statistic and correction for multiple comparisons (FDR<0.05; Benjamini-Hochberg method). Previously validated ductal and myoepithelial cell markers are highlighted in green and red respectively.

### The transcriptional profiles of CD49f^high^/KIT^neg^ and CD49f^low^/KIT^+^ cells are highly conserved across ACCs from different patients

To understand whether CD49f^high^/KIT^neg^ and CD49f^low^/KIT^+^ cells isolated from ACCs of different patients displayed similar gene-expression patterns, we used FACS to sort autologous pairs of cells from all 5 bi-phenotypic PDX lines (ACCX5M1, ACCX6, ACCX14, ACCX22, SGTX6), and then analyzed them by conventional bulk RNA-seq. This allowed us to obtain robust transcriptional profiles of the two populations, in which the representation of genes would not be compromised by technical limitations (i.e., low capture efficiency, gene dropouts) intrinsic to scRNA-seq techniques.

As a first step, we performed a *Principal Component Analysis* (PCA) of the 10 samples included in the RNA-seq dataset **(Figure 1h)**. The analysis revealed that, indeed, the 10 samples grouped tightly into two separate and distinct clusters (5 samples/cluster), exactly corresponding to the two cell phenotypes. Importantly, the analysis also showed that the two clusters separated along the first principal component (PC1), which accounted for a dominant fraction (58%) of the variability in gene-expression levels observed within the dataset. Taken together, these observations indicated that CD49f^high^/KIT^neg^ and CD49f^low^/KIT^+^ cells are defined by systematic differences in transcriptional profiles, which are strongly conserved across tumors originated from different patients. Furthermore, transcriptional differences between the two cell types are greater in magnitude than those attributable to variables which are unique to the individual tumors (e.g., site of origin, repertoire of somatic mutations, genetic background of patients) **(Supplementary Table 1)**.

To understand which genes were differentially expressed between the two cell phenotypes in the most robust and systematic manner, we performed a *differential expression* (DE) analysis using the *DESeq2* package^15^. The analysis identified 643 genes that displayed more than a 2-fold difference in expression levels between the two populations, with high statistical significance (p<0.05, after Benjamini-Hochberg adjustment, **Supplementary Table 2**). We then ranked all differentially expressed genes based on their adjusted p-value, and selected the top 100 for visualization by hierarchical clustering **(Figure 1i).** Among the genes preferentially expressed by CD49f^high^/KIT^neg^ cells were several well-established markers of the myoepithelial lineage (e.g., *ACTA2, MYH11*, *PDPN, TP63*)^16, 17^. Among the genes preferentially expressed by CD49f^low^/KIT^+^ cells were several well-established markers of the ductal and luminal lineage in ectodermal secretory glands (e.g., *ELF5, KIT, SLPI, ANXA8*)^18–21^. Taken together, our observations confirmed that CD49f and KIT represented robust markers for the differential purification of myoepithelial-like (CD49f^high^/KIT^neg^) and ductal-like (CD49f^low^/KIT^+^) populations from human ACCs.

In parallel, we also tested whether CD49f^high^/KIT^neg^ and CD49f^low^/KIT^+^ cells differed in the expression levels of the *MYB* and *NFIB* genes, or *MYB-NFIB* chimeric transcripts, detected using the *STAR-Fusion* software^22^. A majority (50-60%) of human ACCs contain t(6;9) *MYB-NFIB* chromosomal translocations, which cause constitutive upregulation of MYB expression, and are recognized as primary oncogenic drivers in ACC cancerogenesis^5, 6, 23–26^. Although it is established that, when present, the *MYB-NFIB* translocation is found in both myoepithelial-like and ductal-like cells^25^, it remains unknown whether the two cell types actually differ in terms of expression of the corresponding chimeric transcript. Our analysis found no significant difference in the expression level of *MYB* or *NFIB* between the two populations across the 5 analyzed tumors **(Extended Data Fig. 4a-b)**. Out of the 5 PDX models with a bi-phenotypic histology, 3 (SGTX6, ACCX5M1, ACCX14) contained MYB-NFIB fusion transcripts which gave rise to chimeric transcripts, in agreement with previous results from whole genome and Sanger sequencing studies^27, 28^. In these models, fusion transcripts were representative of multiple isoforms, originated from the alternative splicing of the 3’ end (NFIB) of the chimeric gene^23, 24^, and displayed no statistical difference in expression levels between the two populations **(Extended Data Fig. 4c, e)**.

### CD49f^high^/KIT^neg^ cells display a higher tumor-initiating capacity as compared to CD49f^low^/KIT^+^ cells

To understand whether the two populations display functional differences in terms of their capacity to initiate and sustain tumor growth, we compared their tumor-initiating capacity upon prospective xeno-transplantations by *Extreme Limiting Dilution Analysis* (ELDA)^29^. We used FACS to purify autologous pairs of CD49f^high^/KIT^neg^ and CD49f^low^/KIT^+^ cells from bi-phenotypic PDX lines and injected them, side-by-side, in the subcutaneous tissue of NOD/SCID/IL2Rγ^-/-^ (NSG) mice, at progressively decreasing doses (10,000-250 cells/mouse) **(Figure 2a)**. The results showed that the frequency of tumorigenic cells was substantially higher in CD49f^high^/KIT^neg^ as compared to CD49f^low^/KIT^+^ cells. As the dose of injected cells progressively decreased, the frequency of tumors formed by CD49f^low^/KIT^+^ cells was rapidly reduced as compared to the frequency of tumors formed by the CD49f^high^/KIT^neg^ population, until it was eventually lost **(Figure 2d-e, Extended Data Fig. 5)**. To understand whether the higher tumor-initiating frequency observed among CD49f^high^/KIT^neg^ cells could be attributed to higher proliferation rates, we analyzed their cell-cycle distribution and compared it to that of autologous CD49f^low^/KIT^+^ cells, across all 5 bi-phenotypic PDX lines **(Extended Data Fig. 5a-d)**. We found that CD49f^high^/KIT^neg^ cells had a lower proportion of cells in the G2/M phase of the cell cycle compared to CD49f^low^/KIT^+^ cells **(Extended Data Fig. 5g-h)**, in agreement with our scRNA-seq data, which had revealed a sub-group of cells expressing proliferation markers within the ductal-like (*KIT*^+^) population **(Figure 1e, Extended Data Fig. 3)**. Taken together, our results indicated that myoepithelial-like cells represent an aggressive component of the malignant tissue, capable of initiating and sustaining tumor growth, despite their low proliferation kinetics.

**Figure 2:**
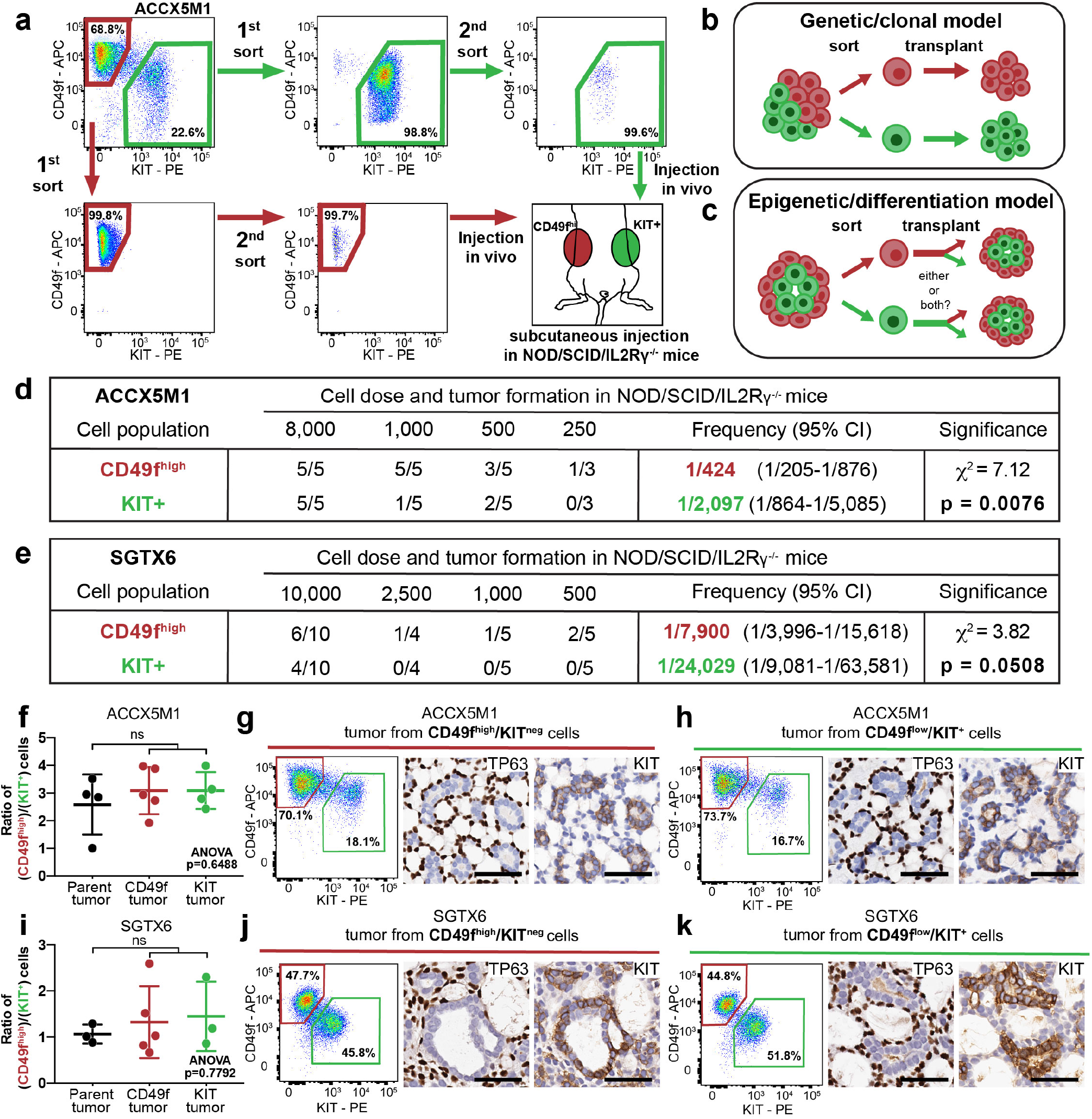
Myoepithelial-like cells (CD49f^high^/KIT^neg^) are highly tumorigenic and act as progenitors of ductal-like (CD49f^low^/KIT^+^) cells. **(a)** Experimental workflow of prospective xeno-transplantation experiments designed to compare CD49f^high^/KIT^neg^ and CD49f^low^/KIT^+^ cells for their tumor-initiating capacity. The two populations were purified in parallel from the same tumor lesion, double-sorted to achieve high purity (>99%) and injected subcutaneously side-by-side, into the opposite flanks of the same animal. **(b)** Predicted outcomes of cell transplantation experiments under a “clonal” model, where each cell type gives rise to a mono-phenotypic outgrowth identical in phenotype to parent cells. **(c)** Predicted outcomes of transplantation under a “differentiation” model, where one or more cell types could serve as a progenitor of the others, in a plastic and dynamic fashion. **(d-e)** *Extreme limiting dilution analysis* (ELDA) of xenotransplantation experiments using paired CD49f^high^/KIT^neg^ and CD49f^low^/KIT^+^ populations sorted from the ACCX5M1 (d) and SGTX6 (e) PDX lines. In both models, the frequency of tumor-initiating cells was higher in CD49f^high^/KIT^neg^ as compared to CD49f^low^/KIT^+^ cells. **(f-k)** Analysis of tumors originated from xeno-transplantation of sorted cells. Analysis by flow cytometry revealed identical ratios of CD49f^high^/KIT^neg^ and CD49f^low^/KIT^+^ cells across parent tumors (black), tumors originated from CD49f^high^/KIT^neg^ cells (red) and tumors originated from CD49f^low^/KIT^+^ cells (green), in both ACCX5M1 **(f-h)** and SGTX6 (i-k) models. Error bars: mean +/- standard deviation. Statistical test: one-way ANOVA with Dunnett’s multiple comparisons test (ns = non-significant, p > 0.05). Analysis by *immunohistochemistry* (IHC) of the same set of tumors confirmed that they all recapitulated a classical cribriform histology, and contained two distinct cell populations, defined by the mutually exclusive expression of myoepithelial-specific (TP63) and ductal-specific (KIT) biomarkers. Identical patterns were observed between parent tumors and tumors originated from sorted cells. Scale bar = 50 µm.

### CD49f^high^/KIT^neg^ cells and CD49f^low^/KIT^+^ cells do not represent distinct genetic clones, but emerge as a result of epigenetic differentiation

We next wanted to understand whether the two cell populations represented different genetic clones, originated from the accumulation of distinct somatic mutations, or different developmental lineages, originated through differentiation processes akin to those that enable stem cells to sustains the homeostasis and regeneration of multi-cellular tissues. We hypothesized that, in a “genetic/clonal” scenario, distinct cell types would maintain their phenotypic identity upon transplantation **(Figure 2b)**. In contrast, in an “epigenetic/differentiation” scenario, one or more cell types could serve as a progenitor of the others, in a plastic and dynamic fashion **(Figure 2c)**. When we analyzed the cell composition of tumors originated from the ELDA experiments, we found that tumors derived from CD49f^high^/KIT^neg^ cells recapitulated the cribriform histology and bi-phenotypic cell composition, with both cell populations (CD49f^high^/KIT^neg^ and CD49f^low^/KIT^+^) present at identical ratios to those observed in the parent lines **(Figure 2f-g, i-j)**. This finding indicated that CD49f^high^/KIT^neg^ cells can differentiate into CD49f^low^/KIT^+^ cells, thus excluding the “genetic/clonal” hypothesis. When we analyzed the limited number of tumors originated from high doses of CD49f^low^/KIT^+^ cells, we also found them to be indistinguishable from parent lines **(Figure 2h, k)**. In the specific case of CD49f^low^/KIT^+^ cells, however, given the higher number of cells required to initiate tumor growth, we could not exclude the possibility that tumors might have arisen from a small cross-contaminations of CD49f^high^/KIT^neg^ cells, despite the high levels of purity achieved by double-sorting.

### CD49f^high^/KIT^neg^ and CD49f^low^/KIT^+^ cells are characterized by differential activation of the *retinoic acid* (RA) signaling pathway

To shed light on the molecular mechanisms involved in regulating the differentiation of CD49f^high^/KIT^neg^ cells into CD49f^low^/KIT^+^ cells, we decided to investigate the RA signaling pathway. RA signaling is critical for proper morphogenesis and differentiation of salivary gland tissues during development^30–32^. Furthermore, recent studies have demonstrated that stimulation of RA signaling can antagonize MYB signaling and slow tumor growth in ACC PDX lines^33, 34^. We first examined whether genes encoding effectors and modulators of RA signaling, including enzymes necessary for RA biosynthesis^30^, RA binding proteins^35, 36^, and RA receptors^37^, were differentially expressed between CD49f^high^/KIT^neg^ cells and CD49f^low^/KIT^+^ cells isolated from PDX lines **(Figure 3a)**. Our analysis revealed that, indeed, most of these genes displayed statistically significant differences in expression levels **(Figure 3b)**. More importantly, when we categorized such genes as potentiators^30, 32, 36^ or attenuators^38, 39^ of RA signaling, we found that potentiators were expressed at higher levels in CD49f^low^/KIT^+^ cells, while attenuators were expressed at higher levels in CD49f^high^/KIT^neg^ cells, in a coordinated and systematic fashion **(Figure 3c)**.

**Figure 3:**
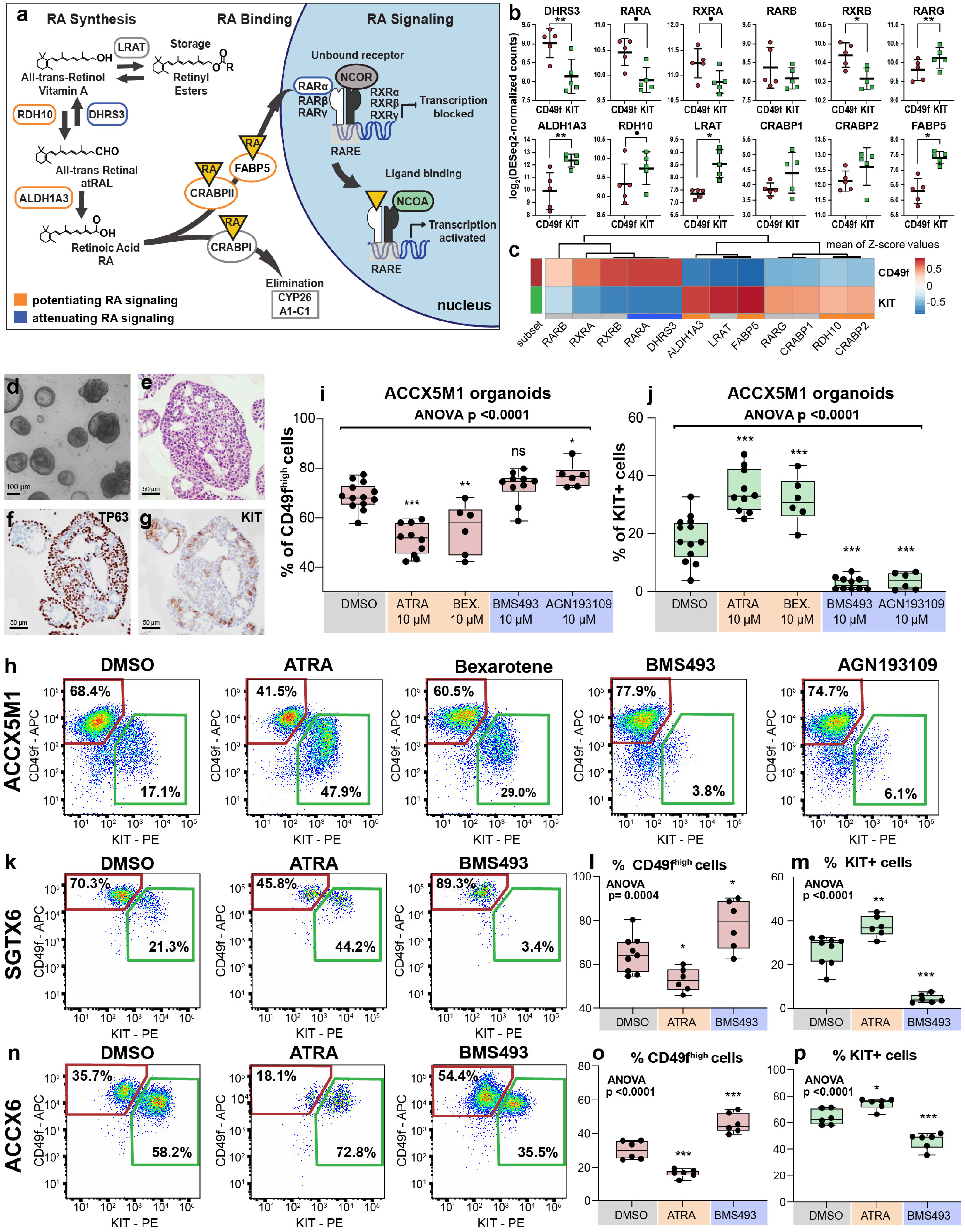
Modulation of *retinoic acid* (RA) signaling can modify the cell composition of human ACCs organoids. **(a)** Schematic modeling of the RA signaling pathway, displaying known potentiators (orange) and attenuators (blue) of RA signaling. **(b)** Comparison of the gene-expression levels of known mediators of RA signaling between myoepithelial-like (CD49f^high^/KIT^neg^; red) and ductal-like (CD49f^low^/KIT^+^; green) cells isolated from bi-phenotypic ACC models. Genes identified as attenuators of RA signaling displayed preferential expression in CD49f^high^/KIT^neg^ cells, while genes identified as potentiators of RA signaling displayed preferential expression in CD49f^low^/KIT^+^ cells, as revealed by RNA-seq. Error bars: mean +/- standard deviation. Statistical test: Student’s t-test for paired samples (• p<0.01, *p<0.05, **p<0.01). **(c)** Heatmap of the mean z-score values for the expression level of genes identified as modulators of RA, as measured across the two ACC populations. **(d-g)** Establishment and phenotypic characterization of *three-dimensional* (3D) organoid cultures from human ACCs. Organoids grow as spherical structures **(d)**, retain cribriform histology **(e)** and contain both myoepithelial-like and ductal-like populations, as visualized by IHC staining for TP63 **(f)** and KIT **(g)**. **(h)** Analysis by flow cytometry of ACC organoids treated for one week with either DMSO (n=13), ATRA (10 µM; n=10), Bexarotene (10 µM; n=6), BMS493 (10 µM; n=10) or AGN193109 (10 µM; n=6). **(i-j)** Comparison of the percentage of CD49f^high^/KIT^neg^ (i) and CD49f^low^/KIT^+^ (j) cells found in organoids following *in vitro* treatment with modulators of RA signaling. Treatment with agonists of RA signaling was associated with an increase in the relative percentage of CD49f^low^/KIT^+^ cells, while treatment with inhibitors of RA signaling was associated with its reduction as compared to DMSO-treated controls. Statistical test: one-way ANOVA with Dunnett’s multiple comparisons test (*p<0.05, **p<0.01, ***p<0.001). **(k-p)** Representative flow plots and quantification of the relative frequency of myoepithelial-like (CD49f^high^/KIT^neg^) and ductal-like (CD49f^low^/KIT^+^) cells in ACC organoids established from SGTX6 **(k-m)** or ACCX6 **(n-p)** PDX lines, treated for one week with either DMSO, ATRA (10 µM) or BMS493 (10 µM), confirming results previously obtained in the ACCX5M1 PDX line. Data correspond to the results of at least two independent experiments (with a minimum of 3 replicates for each treatment condition).

### Activation of the RA signaling pathway induces differentiation of CD49f^high^/KIT^neg^ cells into CD49f^low^/KIT^+^ cells, while inhibition of the RA signaling pathway causes selective death of CD49f^low^/KIT^+^ cells

To formally test whether RA signaling controls cell differentiation in ACCs, we utilized a three-dimensional (3D) *in vitro* organoid culture system. ACC organoid cultures retained the bi-phenotypic composition of PDX models, and thus allowed us to quantify the effects on cell composition caused by pharmacological perturbations of RA signaling **(Figure 3d-g)**^40, 41^. We tested whether activation or inhibition of RA signaling altered the cell composition of ACC organoids, using direct and inverse agonists. We found that agonism of RA signaling with increasing doses of either *all-trans retinoic acid* (ATRA; 1-10 µM) or bexarotene (BEX; 10 µM) led to a progressive increase in the percentage of CD49f^low^/KIT^+^ cells, and a concomitant decrease in the percentage of CD49f^high^/KIT^neg^ cells **(Figure 3h, Extended Data Fig. 6)**. Conversely, inhibition of RA signaling by treatment with increasing doses of either BMS493 (1-10 µM) or AGN193109 (1-10 µM), two pan-RAR inverse agonists, caused a dramatic loss of CD49f^low^/KIT^+^ cells **(Figure 3j, Extended Data Fig. 6).** To ensure the robustness of our findings, we confirmed the effects of this pharmacologic manipulation across three independent PDX lines **(Figure 3h, k-p)**.

To understand the mechanism underpinning the perturbations in cell composition caused by activators and suppressors of RA signaling, we first tested whether ATRA or BMS493 induced selective proliferation of one of the two cell populations. Analysis by IHC of serial sections of 3D organoids revealed no increases in the number of MKI67^+^ cells in either their myoepithelial-like (TP63^+^) or ductal-like (KIT^+^) compartments **(Figure 4a-d, h, l, Extended Data Fig. 7).** Furthermore, analysis by flow cytometry of treated and untreated organoids revealed no changes in cell cycle distribution, indicating that changes in cell composition were not caused by alterations in the proliferation kinetics of either population. **(Extended Data Fig. 7p-s)**. In agreement with our flow cytometry data, however, we observed an increase in KIT^+^ cells in organoids treated with ATRA **(Figure 4g)** and a dramatic loss of KIT^+^ cells in organoids treated with BMS493 **(Figure 4k)**. Remarkably, organoids treated with BMS493 revealed a striking change in morphology, with areas usually occupied by KIT^+^ cells displaying widespread fragmentation of cell nuclei and appearance of amorphous, eosin-rich structures, suggesting a selective cytotoxic effect on ductal-like cells **(Figure 4i, Extended Data Fig. 7)**.

**Figure 4:**
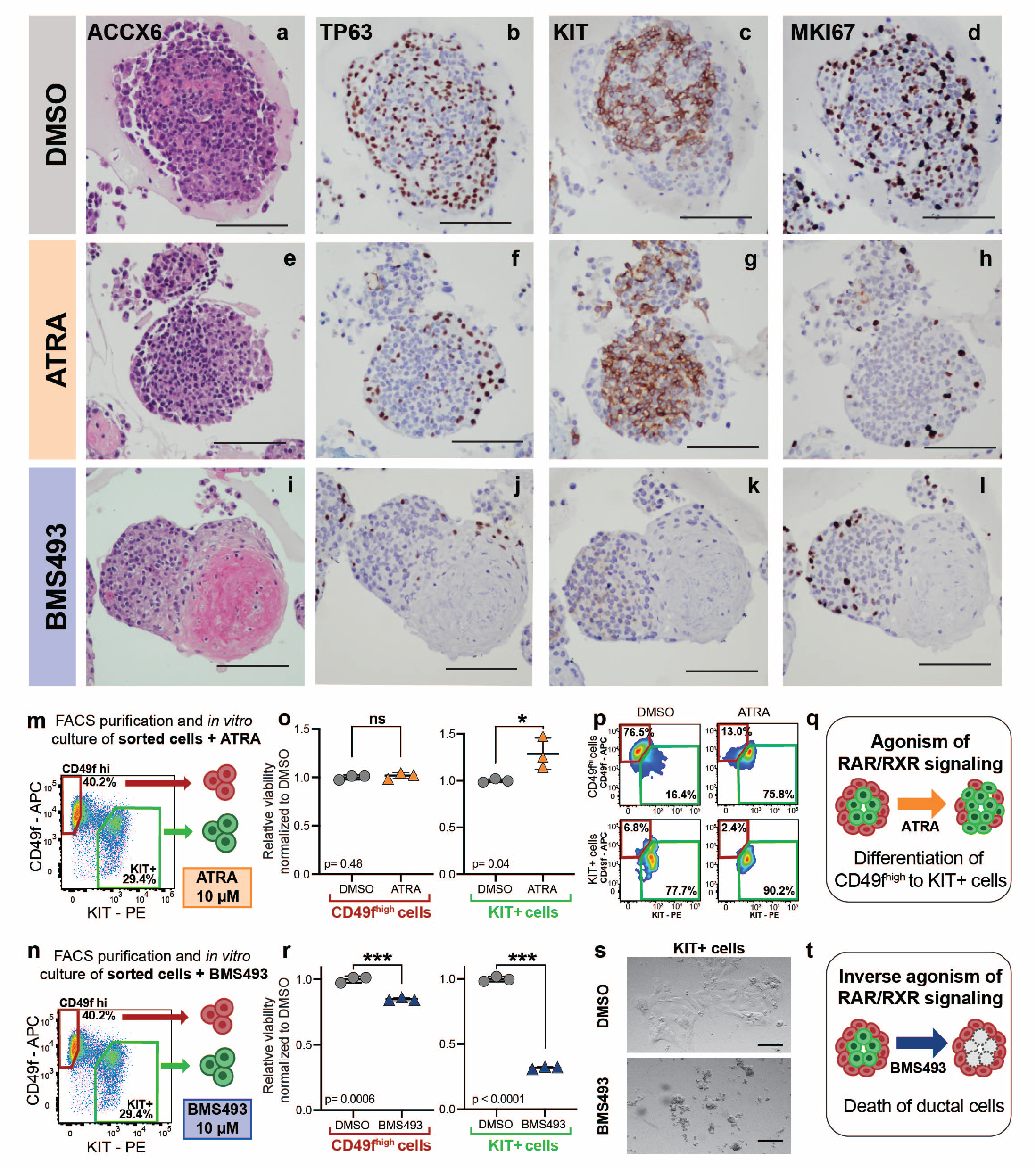
Activation of RA signaling promotes differentiation of myoepithelial-like cells into ductal-like cells, while suppression of RA signaling causes selective death of ductal-like cells. **(a-l)** Organoids established from the ACCX6 PDX line were treated with DMSO, ATRA (10 µM) or BMS493 (10 µM) and subsequently analyzed by IHC. Treatment with ATRA resulted in an expansion of KIT^+^ cells **(g)**, while treatment with BMS493 resulted in a complete loss of KIT expression **(k)**, associated with a dramatic change in the organoids’ morphology, characterized by the appearance of amorphous, eosin-rich deposits at their center **(i)**. Neither ATRA or BMS493 appeared to upregulate MKI67 expression in either cell population **(d, h, l).** Scale bars = 100 µm. **(m-n)** Schematic workflow of prospective experiments aimed at testing the effects of RA signaling on purified populations of myoepithelial-like and ductal-like cells: CD49f^high^/KIT^neg^ and CD49f^low^/KIT^+^ cells were sorted in parallel from the same ACCX5M1 tumors and cultured for one week as 2D monolayers in the presence of either ATRA (10 µM) or BMS493 (10 µM). **(o)** Treatment with ATRA did not reduce the viability of either CD49f^high^/KIT^neg^ or CD49f^low^/KIT^+^ cells (viability assay: alamarBlue; technical replicates: n=3; error bars: mean +/- standard deviation; statistical test: Student’s t-test, two-tailed; ns = non-significant, *p<0.05, **p<0.01, ***p<0.001). **(p)** Treatment with ATRA caused CD49f^high^/KIT^neg^ cells to change phenotype and become undistinguishable from CD49f^low^/KIT^+^ cells by flow cytometry. **(q)** Schematic modeling of the effects of RA agonism on the cell composition of ACC organoids: myoepithelial-like cells are stimulated to differentiate into ductal-like cells. **(r)** Treatment with BMS493 had a minor effect on the viability of CD49f^high^/KIT^neg^ cells, but caused the death of the majority CD49f^low^/KIT^+^ cells. **(s)** Upon visual inspection by conventional microscopy, CD49f^low^/KIT^+^ cells treated with BMS493 appeared fragmented. Scale bar = 100 µm. **(t)** Schematic modeling of the effects of RA inhibition on the cell composition of ACC organoids: ductal-like cells are selectively killed.

To formally dissect the effects of activators and suppressors of RA signaling on the two cell populations, we purified CD49f^high^/KIT^neg^ and CD49f^low^/KIT^+^ cells by FACS and then treated them individually with either ATRA (10 µM) or BMS493 (10 µM) using *in vitro* monolayer cultures^42^ **(Figure 4m, q)**. The experiment revealed that agonism of RA signaling (i.e., treatment with ATRA) did not cause cell death of either CD49f^high^/KIT^neg^ or CD49f^low^/KIT^+^ cells **(Figure 4n)**, but induced a dramatic change in phenotype in CD49f^high^/KIT^neg^ cells, which became almost completely CD49f^low^/KIT^+^ **(Figure 4o)**. This result indicated that, the effects observed in organoid cultures stimulated with RA agonists resulted from differentiation of CD49f^high^/KIT^neg^ cells into CD49f^low^/KIT^+^ cells **(Figure 4p)**. Conversely, the experiment also demonstrated that suppression of RA signaling (i.e., treatment with BMS493) resulted in fragmentation and death of the majority of CD49f^low^/KIT^+^ cells **(Figure 4r-s)**, indicating that the effects observed in organoid cultures following treatment with RA inverse agonists were likely caused by selective cell death of CD49f^low^/KIT^+^ cells **(Figure 4t)**.

### Inverse agonists of RA signaling can be leveraged as anti-tumor agents against human ACCs, with preferential activity against “solid” variants of the disease, which are exclusively composed of CD49f^low^/KIT^+^ cells

We next aimed to elucidate whether the effects observed *in vitro* following treatment with inverse agonists of RA signaling could be leveraged for *in vivo* therapy targeting human ACCs. Because we observed that inverse agonists of RA signaling had selective/preferential cytotoxicity against CD49f^low^/KIT^+^ cells, we hypothesized that tumors containing a large percentage of KIT^+^ cells would represent the most susceptible targets. Although ACCs usually present with a bi-phenotypic histology, over the natural history of the disease a subgroup of ACCs progresses to a “solid” histological pattern, predominantly composed of ductal-like (KIT^+^) cells. Recent studies have shown that ACC progression to solid histology associates with activating mutations in *NOTCH1*^5, 6, 27, 43, 44^, higher proliferation kinetics and a more aggressive clinical course^9, 45, 46^.

To understand whether ACCs with solid histology indeed represented mono-phenotypic expansions of ductal-like cells akin to those found in bi-phenotypic ACC tumors, we analyzed two PDX lines with solid histology and *NOTCH1* activating mutations (ACCX9, ACCX11)^27^. Indeed, analysis of the two PDX lines by FACS revealed that, in both cases, the bulk of the malignant cells consisted of a single population characterized by high expression levels of KIT **(Figure 5a-b)**. Analysis by IHC confirmed that both lines consisted almost exclusively of KIT^+^ cells, and lacked cells expressing the myoepithelial marker TP63, in agreement with previous reports^27^ **(Figure 5c-f)**. We then performed bulk RNA-sequencing of KIT^+^ cells sorted by FACS from both models, in order to obtain a better understanding of their transcriptional profiles. When we performed a PCA analysis, this time combining the two KIT^+^ populations purified from solid ACCs with the 5 paired sets of CD49f^high^/KIT^neg^ and CD49f^low^/KIT^+^ cells isolated from bi-phenotypic ACCs, we observed that, indeed, the KIT^+^ cells from the two solid ACCs clustered with the CD49f^low^/KIT^+^ cells from the bi-phenotypic tumors, indicating that they retained a transcriptional profile characteristic of ductal-like cells **(Figure 5g)**. A similar result was also obtained when we performed a hierarchical clustering of the same 12 populations, using the top 100 genes identified as the most differentially expressed between CD49f^high^/KIT^neg^ and CD49f^low^/KIT^+^ cells **(Figure 5h)**.

**Figure 5:**
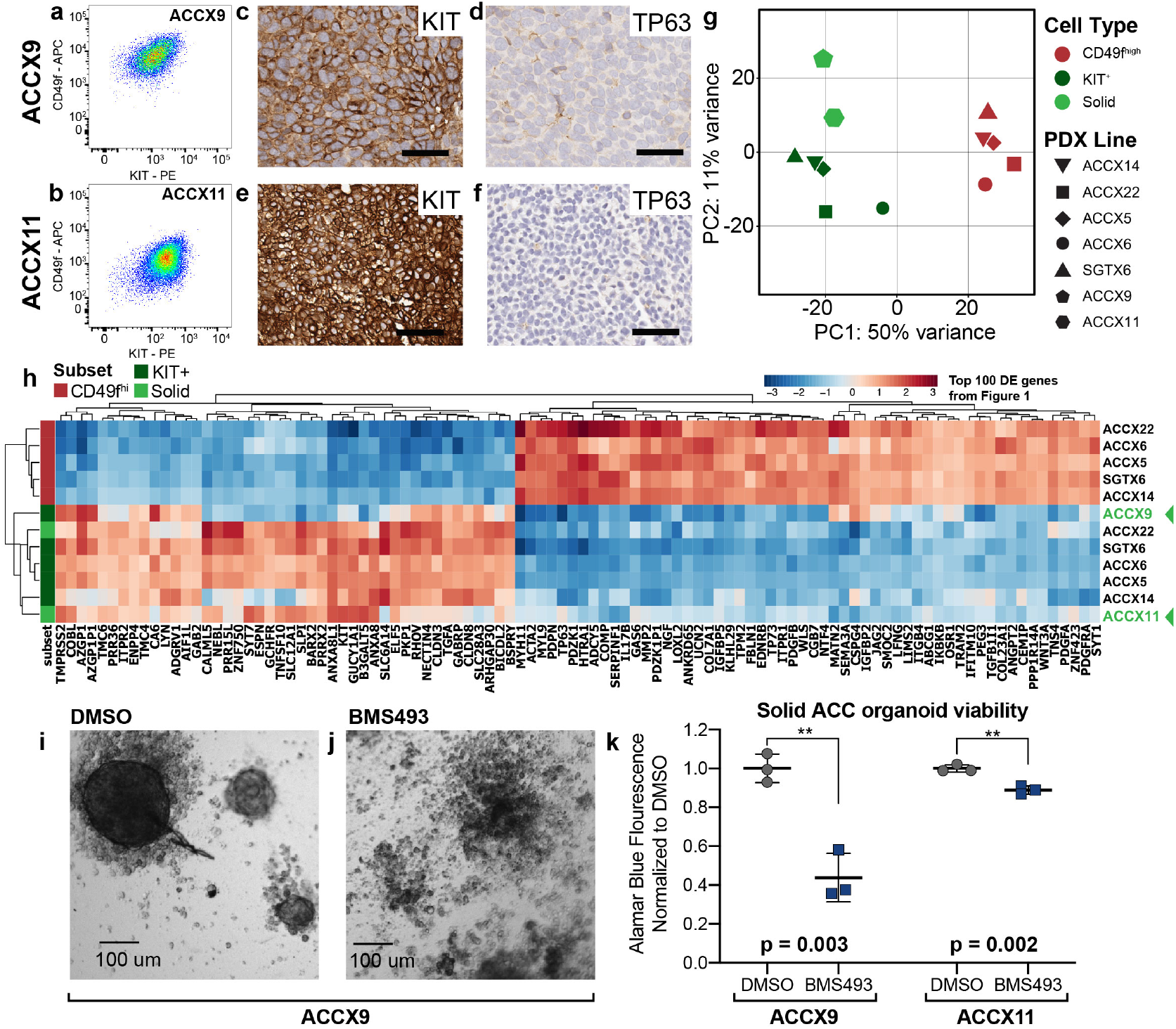
ACCs with solid histology mirror the transcriptional profile of CD49f^low^/KIT^+^ cells from bi-phenotypic ACCs, and are vulnerable to RA inhibition using BMS493. **(a-b)** Analysis by flow cytometry of two PDX lines representative of human ACCs with solid histology (ACCX9, ACCX11) revealing a ductal-like, mono-phenotypic (CD49f^low^/KIT^+^) cell composition. **(c-f)** IHC analysis of KIT and TP63 expression ACCX9 and ACCX11, showing ubiquitous expression of the ductal-specific marker KIT **(c,e)** and loss of the myoepithelial-specific marker TP63 **(d,f)**. Scale bars = 50 µm. **(g)** *Principal component analysis* (PCA) of RNA-seq data from human ACCs, in which results from two PDX lines with solid histology (ACCX9, ACCX11) are combined with 5 autologous pairs of CD49f^high^/KIT^neg^ and CD49f^low^/KIT^+^ cells populations from bi-phenotypic PDX lines (ACCX5M1, ACCX6, ACCX14, ACCX22, SGTX6). **(h)** Hierarchical clustering of RNA-sequencing data from human ACCs, based on the expression levels of the top 100 genes identified as differentially expressed between CD49f^high^/KIT^neg^ and CD49f^low^/KIT^+^ cells. Solid ACCs cluster with CD49f^low^/KIT^+^ cells from bi-phenotypic tumors, irrespective of the method used to analyze their transcriptional profile. **(i-j)** Upon visual inspection by conventional microscopy, ACCX9 organoids cultured for one week in the presence of BMS493 (10 µM) display widespread cell fragmentation, in contrast to organoids cultured with DMSO alone. **(k)** Quantification of organoid viability using the alamarBlue HS assay, confirming the cytotoxic activity of BMS493 (10 µM) against PDX lines with solid histology (ACCX9, ACCX11). Error bars: mean +/- standard deviation; p-values: two-tailed Student’s t-tests; n=3 replicates/condition.

We then wanted to understand whether, in addition to sharing transcriptional similarities with the CD49f^low^/KIT^+^ cells of bi-phenotypic ACCs, KIT^+^ cells from solid ACCs also shared a vulnerability to pharmacological suppression of RA signaling. We generated organoid cultures from both PDX lines with solid histology, and then treated them *in vitro* with BMS493 (10 µM). After one week of culture, organoids treated with BMS493 began to lose structural integrity, appearing fragmented and apoptotic **(Figure 5j)**. In both cases, treatment with BMS493 resulted in a statistically significant decrease in cell viability **(Figure 5k)**.

As a final step, we wanted to test whether an inverse agonist of RA signaling (BMS493) could be leveraged for the *in vivo* treatment of human ACCs with either solid (mono-phenotypic) or cribriform (bi-phenotypic) histology. We engrafted three PDX lines, two representative of solid ACCs (ACCX9, ACCX11) and one representative of a cribriform ACC (ACCX5M1), in NOD/SCID/IL2Rγ^-/-^ (NSG) mice. We then let tumors grow up to a volume of approximately 0.4 cm^3^, and then treated tumor-bearing mice with BMS493 (40 mg/kg, i.p.) using a slightly more intense regimen for the cribriform models (4 times/week x 3 weeks) as compared to the solid models (3 times/week x 3 weeks), under the assumption that the cribriform model would be more resistant to the treatment. As expected, treatment with BMS493 was associated with side-effects reminiscent of those previously described as secondary to vitamin A deficiency in rodents (e.g., encrusted eyelids, rough coat, scaling of skin)^47^, which began to manifest during the second week of treatment and worsened during the third. Overall, out of 18 tumor-bearing animals treated with BMS493, 33% (n=6/18) experienced reductions in tumor volume **(Extended Data Fig. 8)**. Four animals (4/18 22%) had to be prematurely euthanized due to a sudden deterioration in general health; in three of these animals, euthanasia followed tumor shrinkage, suggesting a possible acute complication due to widespread tumor lysis (**Extended Data Fig. 8**). Overall, treatment with BMS493 was associated with a statistically significant reduction in the growth kinetics of engrafted tumors across all three tested models, even after removal of the four mice undergoing premature euthanasia **(Figure 6c, e, h, Extended Data Fig. 8)**. Taken together, our data revealed that BMS493 has robust anti-tumor activity against human ACCs, both *in vitro* and *in vivo*, and indicate that inverse agonist of RA signaling are deserving of further study as a new class of anti-tumor agents for the clinical treatment of salivary gland malignancies.

**Figure 6:**
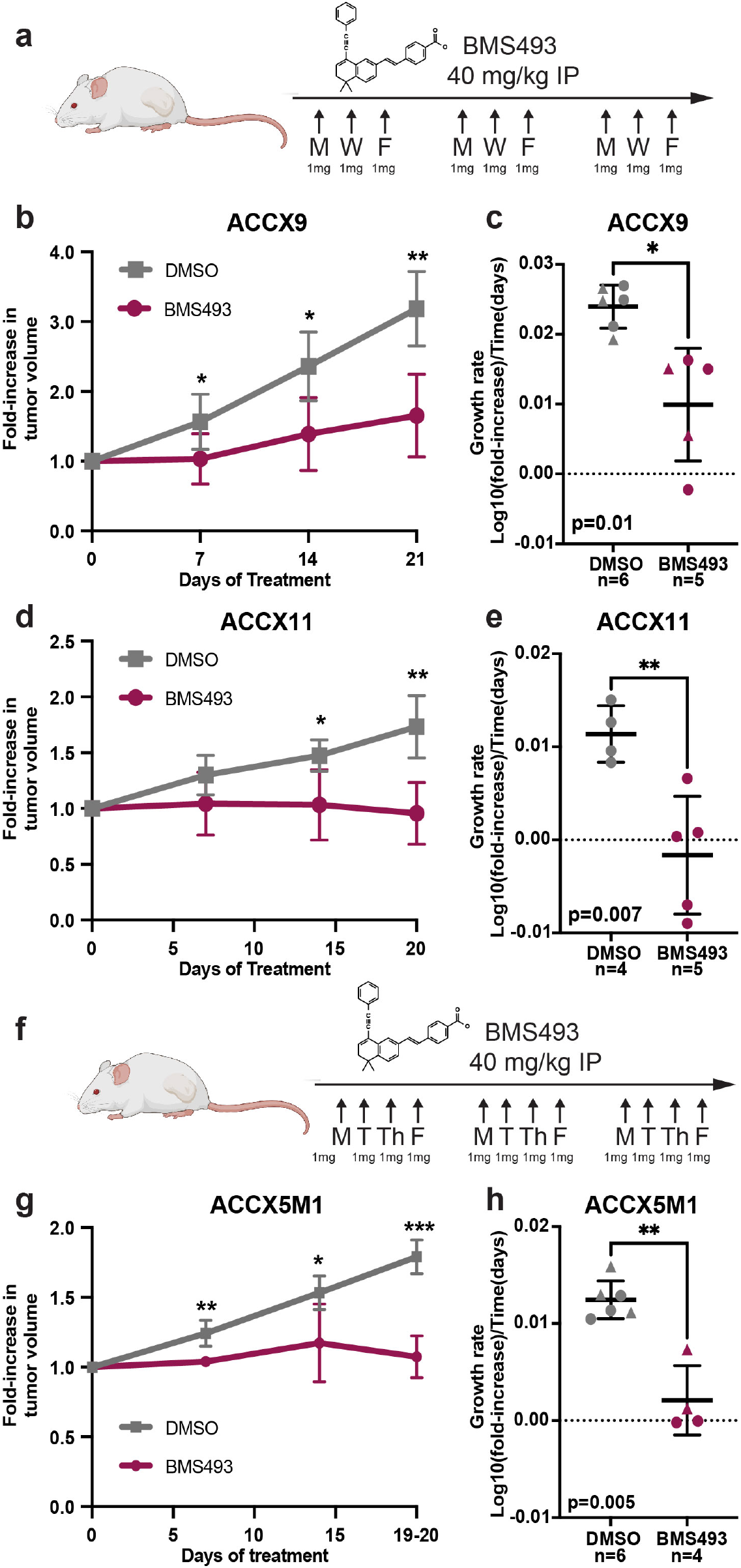
*In vivo* anti-tumor activity of BMS493. **(a**) Schematic of the BMS493 dosing regimen utilized for the *in vivo* treatment of solid ACC models (40 mg/kg doses, i.p., 3 times/week x 3 weeks). Schematic was created with BioRender.com. **(b-e)** Comparison of tumor growth kinetics, expressed as either fold-increases in mean tumor volumes or growth rates, between mice treated with BMS493 and mice treated with the drug’s vehicle alone (DMSO), following subcutaneous engraftment with solid ACC models (ACCX9: **b-c**; ACCX11: **d-e**). **(f)** Schematic of the BMS493 dosing regimen utilized for the *in vivo* treatment of a bi-phenotypic ACC model (40 mg/kg doses, i.p., 4 times/week x 3 weeks). **(g-h)** Comparison of tumor growth kinetics, expressed as either fold-increases in tumor volumes or growth rates, between mice treated with BMS493 and mice treated with the drug’s vehicle alone (DMSO), following subcutaneous engraftment with a bi-phenotypic ACC model (ACCX5M1). Differences between mean fold-increases in tumor volume were tested for statistical significance using a two-tailed Student’s t-test (ns = non-significant, *p<0.05, **p<0.01, ***p<0.001). Differences between mean growth rates were tested for statistical significance using a two-tailed Welch’s t-test. Growth rates were calculated assuming exponential kinetics. Error bars: mean +/- standard deviation.

## DISCUSSION

The results of our study reveal that the two major sub-types of malignant cells (myoepithelial-like and ductal-like), which co-exist within human ACCs, can be differentially purified by FACS using two surface markers (CD49f, KIT). We find that these two markers are differentially expressed between ACC cell populations in a robust and reproducible manner, and across multiple tumor models. This technical advancement has important implications, as it enables, for the first time, experiments that are aimed at comparing the functional properties of the two populations in *“*side-by-side*”* prospective assays, including experiments designed to elucidate their developmental relationships.

The results of our gene-expression studies, conducted on autologous pairs of cell populations sorted in parallel from the same tumors, indicate that the two populations are defined by transcriptional signatures that are closely recapitulated across different patients. The two cell-specific transcriptional programs appear to represent one of the most dominant sources of intra-tumor heterogeneity, irrespective of each tumor’s mutational repertoire, its site of primary origin, or the patient’s sex. The results of our xeno-transplantation studies in immune-deficient mice, designed to compare the tumorigenic capacity of the two cell populations, show that the two populations do not represent distinct genetic clones (i.e., they do not originate from the divergent accumulation of independent somatic mutations), but distinct epigenetic lineages (i.e., different cell types originated through the dynamic and systematic activation of *“*hard-wired*”* developmental programs, akin to those that enable stem cell populations to support the homeostatic turnover and/or regeneration of normal tissues)^10^. In agreement with these observations, we also find that the relative proportion of myoepithelial-like and ductal-like cells within ACC tumors represents a stable, intrinsic feature of each PDX line, as each PDX model maintains the same relative frequencies over serial passaging, and most notably, in tumors originated from purified cell populations. From a translational point of view, one the most important findings of our study is the observation that, despite their apparent low proliferation rates, myoepithelial-like cells (CD49f^high^/KIT^neg^), represent a highly tumorigenic component of human ACCs, capable of supporting the formation of tumors when transplanted in immune-deficient mice, and acting as a progenitors of the more proliferative ductal-like cells (CD49f^low^/KIT^+^). Traditionally, in tumors originated from secretory glands of ectodermal origin (e.g. breast cancer), myoepithelial-like components are considered to represent a relatively indolent subset of malignant cells, often used as a biomarker of tumors with a more benign clinical behavior, and postulated to exert a suppressive effect on metastatic spread^48, 49^. Our findings caution against this interpretation in the case of the majority of human ACCs, especially those originated from salivary glands. Indeed, our findings indicate that, in order to be curative, treatment strategies for ACCs will likely need to eradicate both myoepithelial-like and ductal-like cell populations.

From a more mechanistic point of view, our study also demonstrates that, in human ACCs, the capacity of myoepithelial-like cells to differentiate into ductal-like cells is controlled by RA signaling. Recent studies have identified RA signaling as a critical pathway in the biology of human ACCs, demonstrating that RA agonists, such as ATRA, have powerful anti-proliferative activity in PDX models^33, 34^. A recent clinical trial, however, showed that treatment with ATRA was associated with limited clinical benefit in ACC patients^50^. Our *in vitro* studies, using 3D organoids and purified cell preparations of both myoepithelial-like and ductal-like cells, provide novel mechanistic insights into the anti-tumor activity of retinoids in ACCs, and a possible explanation for the conflicting results between pre-clinical and clinical data. We find that activation of RA signaling induces differentiation of myoepithelial-like cells into ductal-like cells, without compromising their viability, in agreement with the well-known function of ATRA as a differentiation agent across multiple tissues^51^. We therefore hypothesize that, in a clinical setting, the therapeutic benefit of ATRA might be short-lived, because of the transient and dynamic nature of the perturbations that it is predicted to induce in the cell composition of ACC tumors.

Perhaps more importantly, we found that inhibition of RA signaling induces selective cell death of ductal-like cells. This finding provides an opportunity for the development of novel therapeutic agents against ACCs, especially in tumors with solid histology, which are characterized by a mono-phenotypic expansion of ductal-like cells. These tumors usually originate during the progression of human ACCs, with kinetics that are reminiscent of the “blast crisis” phase of human *chronic myelogenous leukemias* (CMLs), in which a population of more differentiated, yet highly proliferating myeloid progenitors becomes dominant, as a result of mutations that aberrantly activate the pathways that confer self-renewal to normal hematopoietic stem cells^10, 52^. Our pre-clinical data show that, in solid ACCs, inhibition of RA signaling using inverse agonists (e.g., BMS493) can have profound and robust anti-tumor activity. Unlike agonists of RA signaling, which have been extensively explored as anti-tumor agents in a number of human malignancies^53–55^ and have known and manageable associated toxicities^56^, inverse agonists of RA signaling, to our knowledge, have not yet been evaluated for clinical use. Because RA signaling is critical to the biology of many tissues, the anti-tumor benefit of RA antagonists will have to be weighted carefully against any potential toxicities, based on the results of future toxicology and pharmacodynamics studies.

## METHODS

### Data Availability Statement

The RNA-seq datasets generated as part of this study are available in the dbGAP repository (https://www.ncbi.nlm.nih.gov/gap), under accession number: **phs002764.v1.p1**.

### Code availability statement

All computer software used in this study is either publicly or commercially available, and is described in detail under each section of the methods describing the experimental procedures that involved its use.

The software used for single-cell RNA-seq analysis includes:

- *cellranger* (version 3.1.0), available at: https://support.10xgenomics.com/single-cell-gene-expression/software/downloads/latest;
- *Randomly*^12^, available at: https://rabadan.c2b2.columbia.edu/html/randomly;
- *Scanpy*, available at: https://scanpy.readthedocs.io/en/stable;

The software used for bulk RNA-seq analysis includes:

- bcl2fastq2 (version 2.20), available at: https://support.illumina.com/downloads/bcl2fastq-conversion-software-v2-20.html;
- kallisto (version 0.44.0), available at: https://github.com/pachterlab/kallisto;
- DESeq2^15^ (1.28.1), available at: https://bioconductor.org/packages/release/bioc/html/DESeq2.html;
- R (version 4.0.1) https://www.r-project.org/;
- R packages: ggplot2 (version 3.3.3), pheatmap (version 1.0.12), RColorBrewer (version 1.1-2), dplyr (version 1.0.5), tidyverse (version 1.3.0), DEGreport (version 1.24.1), tibble (version 3.1.0), reshape2 (version 1.4.4), GOstats (version 2.54.0), and sva (version 3.36.0, containing ComBat-seq^57^, https://www.bioconductor.org/packages/release/bioc/html/sva.html);
- STAR-fusion^22^ (version 1.7.0), available at: https://github.com/STAR-Fusion/STAR-Fusion

The software used for *Extreme Limiting Dilution Analysis* (ELDA)^29^ is publicly available at: http://bioinf.wehi.edu.au/software/elda. The acquisition and contrast-enhancement of images representative of tissues analyzed by *immunohistochemistry* (IHC) was performed using the *QuPath* software (https://qupath.github.io/, version 0.2.3) and *Adobe Photoshop* (version 22.5.0).

### Patient-derived xenograft (PDX) lines

PDX lines established from 7 independent human ACCs (ACCX5M1, ACCX14, ACCX22, SGTX6, ACCX6, ACCX9, ACCX11) were obtained from the *Adenoid Cystic Carcinomas Registry* (ACCR) at the University of Virginia^13^. Tumor tissues were propagated in NOD/SCID/IL2Rγ^-/-^ (NSG) mice (The Jackson Laboratory; stock #005557) by sub-cutaneous xenotransplantation of solid fragments, following previously published procedures^14^. Animal experiments were approved by the *Institutional Animal Care and Use Committee* (IACUC) of Columbia University, under protocol numbers: AC-AAAL7751, AC-AAAW1466, AC-AABM9553.

### Immunohistochemistry (IHC)

Freshly isolated tissue-specimens were washed in *Dulbecco’s Phosphate Buffer Solution* (DPBS) and fixed overnight (12-18 hours) in a 10% formalin solution (Sigma, HT501320). *Formalin-fixed, paraffin-embedded* (FFPE) tissue-blocks were stained using the *Benchmark Ultra* automated platform (Ventana) with the UltraView DAB detection kit. *Heat-induced epitope retrieval* (HIER) was performed using Cell Conditioning 1 (pH 7.3) solution. Slides were stained (32 minutes) with the following primary antibodies: anti-human TP63 [clone 4A4; Ventana], anti-human KIT [clone YR145; Cell Marque], anti-human MKI67 [clone 30-9; Ventana]. Stained slides were imaged using a digital scanner (Leica SCN400), and regions of interest were captured using the *QuPath* software (https://qupath.github.io/, version 0.2.3). Image brightness and contrast were adjusted using *Adobe Photoshop* (version 22.5.0). Adjustments were applied uniformly to the entire image.

### Tissue dissociation and preparation of single-cell suspensions

ACC tumors were harvested from NSG mice and washed with cold (4°C) DPBS. Tissue was cut into small pieces with surgical scissors, followed by thorough mechanical mincing with a razor blade. The resulting tissue fragments were resuspended in “disaggregation medium”, consisting of: RPMI-1640 medium (Sigma, R8758) supplemented with 2 mM L-alanyl-L-glutamine (Corning 25-015-CI), 100 U/mL penicillin and 100 µg/mL streptomycin (Sigma, P4333), 1x Antibiotic Antimycotic Solution (Corning 30-004-Cl), 20 mM HEPES (Corning, 25-060-CI), 1 mM sodium pyruvate (Gibco, 11360070), 100 units/ml Hyaluronidase (Worthington, LS002592), 100 units/ml DNase-I (Worthington, LS002139), and 200 units/ml Collagenase-III (Worthington, LS004183). Tissue fragments were then incubated at 37°C for two hours, with pipetting every 10-15 minutes to promote cell dissociation. The resulting cell suspension was then serially filtered through 70-µm and 40-µm nylon meshes, in order to remove undigested tissue fragments and cell clumps. *Red blood cells* (RBCs) were removed by osmotic lysis, achieved by incubating the cell-suspension (5 minutes, on ice) in a hypotonic buffer (155 mM ammonium chloride, 0.01 M Tris-HCl; Red Blood Cell Lysing Buffer Hybri-Max; Sigma, R7757). Dissociated single cells were then spun at 1,500 RPM for 5 minutes, and re-suspended by gentle pipetting in “flow cytometry buffer” (FCB) solution, consisting of: 1x Hank’s Balanced Salt Solution (HBSS, Sigma H6648) with 2% heat-inactivated adult bovine serum (Sigma, B9433), 20 mM HEPES (Corning, 25-060-Cl), 5 mM EDTA (Sigma, 3690), 1 mM sodium pyruvate (Gibco, 11360-070), 100 U/ml penicillin and 100 µg/ml streptomycin (Sigma, P4333), and 1x Antibiotic Antimycotic solution (Corning 30-004-Cl).

### Staining of single-cell suspensions with fluorescence-conjugated monoclonal antibodies

To prevent unspecific binding of antibodies, cells were incubated with human IgGs (Innovative Research, VN00089472, 5 mg/ml) in FCB, on ice (4°C) for 15 minutes. Cells were then washed with FCB, and stained (15 minutes, 4°C) with various monoclonal antibodies, each used at a dilution determined by individual titration experiments. Antibodies used for removal of mouse stromal cells included: anti-mouse H-2K^d^-biotin (BioLegend, 116604, clone SF1.1, dilution 1:20) and anti-mouse Cd45-PE/Cyanine5 (BioLegend 103110, clone 30-F11, dilution 1:100). Biotin-conjugated antibodies were visualized by secondary staining with streptavidin PE/Cyanine5 (BioLegend 405205, dilution 1:200). Antibodies used for staining of human tumor cells included: anti-human EpCAM-FITC (BioLegend 324204, clone 9C4, dilution 1:30), anti-human/mouse CD49f-APC (BioLegend 313616, clone GoH3, dilution 1:40) and anti-human KIT-PE (BioLegend 313204, clone 104D2, dilution 1:50). After staining, cells were washed with 1 mL FCB to remove excess unbound antibodies and resuspended in FCB containing DAPI (Invitrogen D3571, dilution 1:10,000).

### Fluorescence-activated cell sorting (FACS)

Single-cell suspensions were either analyzed using a high-parameter flow cytometer (LSRFortessa; Becton Dickinson) or used as starting material to purify selected sub-populations using a cell-sorter (FACSAria; Becton Dickinson). In experiments performed using the LSRFortessa, cell doublets were eliminated using a sequential gating strategy, based on *forward-scatter area* vs. *forward-scatter width* (FSC-A vs. FSC-W) and *side-scatter area* vs. *side-scatter width* (SSC-A vs. SSC-W) profiles. In experiments performed using the FACSAria, cell doublets were eliminated using a similar strategy, with sequential gating based on *forward-scatter area* vs. *forward-scatter height* (FSC-A vs. FSC-H) and *side-scatter area* vs. *side-scatter height* (SSC-A vs. SSC-H) profiles **(Extended Data Fig. 1)**. Dead cells and cells of murine origin (i.e., cells expressing mouse stromal markers, such as H-2K^d^ and Cd45) were eliminated by exclusion of DAPI^+^ and PE/Cyanine5^+^ cells, respectively **(Extended Data Fig. 1)**. Data was acquired using the FACSDiva software (Becton Dickinson) and analyzed using *FlowJo* (Becton Dickinson, version 10.7.1).

### Cell-cycle analysis

Cells were stained with monoclonal antibodies aimed at enabling the differential visualization of CD49f^high^/KIT^neg^ and CD49f^low^/KIT^+^ cells, and then fixed using the BD Cytofix/Cytoperm Fixation/Permeabilization Kit (BD Biosciences, 554722; 20 minutes at 4°C). Cells were then washed twice in 1x BD Perm/Wash buffer, resuspended in Perm/Wash buffer with DAPI (1:1000) and incubated in the dark (30 minutes, 4°C). Finally, cells were washed twice with 1x Perm/Wash buffer, resuspended in FCB and analyzed on the LSRFortessa flow cytometer.

### Single-cell RNA-sequencing (scRNA-seq)

Live, human cancer cells (DAPI^neg^, H-2Kd^neg^, Cd45^neg^, EpCAM^+^) were purified by FACS from a xenograft (ACCX22) and single-cell libraries were prepared using the Chromium Single Cell 3’ Solution (10x Genomics) with the Single Cell 3’ v3 chemistry, following the manufacturer’s instructions. RNA-sequencing was performed on the NovaSeq 6000 (Illumina) at the JP Sulzberger Columbia Genome Center. Sequencing reads were mapped to human transcriptome GRCh38-3.0.0 and analyzed with the *cellranger* pipeline (version 3.1.0; 10x Genomics).

### Single-cell transcriptomics analysis and clustering

Data output from *cellranger* was filtered to retain only cells with sufficient library complexity (i.e., cells expressing >500 genes), yielding a final dataset composed of 3,533 cells. The final expression matrix was normalized based on log_2_(1+TPM) (Transcripts Per Million) values. Gene-expression data was then processed using the *Randomly*^12^ algorithm (Python package available at: https://rabadan.c2b2.columbia.edu/html/randomly/). This algorithm is based on the statistical principles of *Random Matrix Theory* (RMT), and can be used to distinguish between biological and technical sources of signal in single-cell datasets^12, 58^. The algorithm is based on the observation that single-cell data contains a threefold structure composed of: 1) a sparsity-induced, non-biological signal; 2) a random matrix and 3) a true biological signal. The algorithm works by 1) removing genes that induce spurious signal due to sparsity; 2) detecting the part of the signal that corresponds to a random matrix, by identifying eigenvalues that lie outside the *Marchenko-Pastur* (MP) distribution, along their corresponding eigenvectors (**Extended Data Fig. 2a**); and 3) selecting the genes mostly responsible for biological signal. Genes responsible for biological signal are selected by projecting the original expression matrix into the signal-like eigenvectors and the noise-like eigenvectors and performing a chi-squared test for the normalized sample variance which allows for the identification of signal-like genes, based on a specified *False Discovery Rate* (FDR). In the specific case of our experiment on cells purified from the PDX line ACCX22, the *Randomly* pipeline identified 47 informative eigenvectors and approximately 5,500 informative genes, with an FDR ≤ 0.001 **(Extended Data Fig. 2b-c)**. The final output produced by *Randomly* corresponds to the latent space, which is then analyzed to identify cell populations, using synergistic machine-learning techniques for cell clustering. In our experiment, clustering was performed using the Leiden algorithm as implemented in the code by Wolf et al.^59^ (https://scanpy.readthedocs.io/en/stable). The optimal number of clusters was selected based on their mean silhouette score^60^. More precisely, we performed a set of clusterings using different Leiden resolutions, then computed the mean silhouette score for each case, and finally selected the clustering solution with the highest mean silhouette score **(Extended Data Fig. 2d)**.

### Visualization of scRNA-seq results and *differential expression* (DE) analysis

Dimensionality reduction and *Uniform Manifold Approximation and Projection* (UMAP) representation of the scRNA-seq data were performed using the visualization functions of the *Randomly* package, with default parameters. Genes with differential expression between the myoepithelial-like cell-population (Cluster 1) and the ductal-like cell population (Cluster 2) were identified based on the following two-step approach: 1) of all the genes identified as informative based on their projection in the 47 eigenvectors corresponding to biological signal, those displaying preferential expression in one of the two cell populations (i.e., expressed in at least 60% of cells in one population, and no more than 20% of cells the other) were selected for analysis; 2) genes selected for analysis were tested for statistical significance of their difference in mean expression levels between the two populations (Student’s t-test, two-tailed). Those identified as displaying the most robust difference after adjustment for multiple comparisons (FDR <0.001; Benjamini-Hochberg correction) were defined as differentially expressed. Finally, for each of the genes identified as differentially expressed, the distribution of expression levels in each of the two populations was visualized with a dot-plot, whereby the size of each dot represents the percentage of cells expressing the gene in a specific cluster, and the color of the dot represents the gene’s average level of expression within the same cluster.

### RNA-sequencing

Paired sets of CD49f^high^/KIT^neg^ and CD49f^low^/KIT^+^ cell populations were sorted in parallel using a FACSAria (Becton Dickinson). Sorted cells were washed with 1 ml of cold DPBS, pelleted, snap frozen in liquid nitrogen, and stored at −80°C until RNA extraction. RNA was extracted using the NucleoSpin® RNA XS Kit (Takara Bio, 740902.50). RNA quality and abundance were analyzed using a *Bioanalyzer 2100* (Agilent RNA Pico Assay). All samples used for RNA-seq experiments were deemed of satisfactory abundance (total amount >100 ng) and structural integrity (RNA integrity number [RIN] >7). RNA-seq libraries were prepared using the TruSeq stranded mRNA kit (Illumina), after poly-A pull-down to enrich for mRNAs, following the manufacturer’s instructions. RNA-seq experiments were performed in three batches: Batch 1 included paired CD49f^high^/KIT^neg^ and CD49f^low^/KIT^+^ populations from ACCX5M1 and SGTX6, Batch 2 included paired CD49f^high^/KIT^neg^ and CD49f^low^/KIT^+^ populations from ACCX14, and Batch 3 included paired CD49f^high^/KIT^neg^ and CD49f^low^/KIT^+^ populations from ACCX6 and ACCX22, and KIT^+^ cells from ACCX9 and ACCX11. RNA-seq libraries were sequenced using either the Illumina HiSeq 4000 (Batch 1) or the Illumina NovaSeq 6000 (Batch 2, Batch 3) platforms, available at the JP Sulzberger Columbia Genome Center. Batch 1 samples were sequenced to a depth of 30 million reads with single-end 100 base pair read lengths, while Batches 2 and 3 samples were sequenced to a depth of 20 million reads with paired-end 100 base pair read lengths. The *real-time analysis* (RTA) software (Illumina) was used to generate *binary base calls* (BCLs), and the *bcl2fastq2* software (version 2.20) was used for converting BCLs into FASTQ format, coupled with adaptor trimming. Sequencing reads were mapped to human transcriptome GRCh38 using the *kallisto* software (version 0.44.0).

### Bioinformatic analysis of bulk RNA-seq data

Analysis of bulk RNA-seq data was performed in R (version 4.0.1). Data was normalized for batch effects using *ComBat-seq*^57^, and genes differentially expressed between CD49f^high^/KIT^neg^ and CD49f^low^/KIT^+^ cells were identified using the *DESeq2* package^15^. For downstream analyses, including visualization of gene expression levels, clustering, and *principal component analysis* (PCA), gene expression data was transformed using the *rlog* function, which transforms data to the log2 scale, after normalization of read counts with respect to library size. PCA was performed using the *plotPCA* function with default parameter for the number of top genes to use for principal components (i.e., 500 genes). Differentially expressed genes were defined as those displaying a more than two-fold difference in mean expression levels between the two populations (log_2_ fold-change > 1) that was considered to be statistically robust after ranking based on the Wald statistic and correction for multiple comparisons (FDR<0.05; Benjamini-Hochberg method). The variance in gene-expression levels across independent samples was visualized using heatmaps, generated using the *pheatmap* function, with scaling performed by mean-centering the gene expression values for each gene. Heatmaps were organized by hierarchical clustering of both genes and samples.

### Identification and quantification of *MYB-NFIB* chimeric transcripts in RNA-seq data

RNA-seq datasets were analyzed for the presence of *MYB-NFIB* chimeric transcripts, as well as for differences in the relative representation of splicing isoforms, using the *STAR-fusion* software (v1.7.0)^22^, after mapping sequencing results (FASTQ files) to human reference genome GRCh37. Differences in the expression levels of chimeric transcripts, expressed as *fusion fragments per million* (FFPM), were tested for statistical significance using a Student’s t-test for paired samples (two-tailed).

### Statistical analysis

The distribution of experimental data was visualized using either dot-plots or box-plots, generated in GraphPad Prism (version 8) or R (version 4.0.1). In both dot-plots and box-plots, all experimental replicates were reported as individual data-points. In the case of dot-plots, horizontal bars correspond to means and error bars correspond to the standard deviation. In the case of box-plots, boxes correspond to the range of values between the upper and lower quartiles of the data distribution, horizontal bars correspond to the median, and whiskers correspond to the minimum and maximum data-points. The statistical significance of observed differences was evaluated using a variety of tests, chosen on a case-by-case basis, depending on experimental assumptions and data distributions. For each of the reported experiments, the corresponding statistical tests are described both under the corresponding paragraph of the Methods section and the corresponding figure legends. Tests included one-way ANOVA, Student’s t-test (two-tailed; for either paired or unpaired samples, depending on experimental design) and Welch’s t-test (two-tailed; when sample variances were deemed to be unequal based on an F-test). In the specific case of high-throughput experiments involving the simultaneous measurement of the expression levels of thousands of genes (e.g., scRNA-seq, RNA-seq), the identification of differentially expressed genes was based on *false-discovery rates* (FDR) calculated using the Benjamini-Hochberg method, in order to adjust for multiple comparisons.

### Tumorigenicity and ELDA

Paired sets of autologous CD49f^high^/KIT^neg^ and CD49f^low^/KIT^+^ cell populations were double-sorted by FACS starting from the same tumor specimens, using a cell-sorter equipped for 2-way parallel purification (FACSAria, Becton Dickinson) as described above **(Extended Data Fig. 2b-c)**. Double-sorting consisted in two sequential rounds of sorting, whereby, after the first sort, cells were spun down, resuspended in 0.5 mL of fresh FCB with DAPI, and then sorted a second time, using identical gates. Cells were assessed for purity and viability after the second sort, resuspended in fresh FCB and counted using a hemocytometer. Cells were then aliquoted at various doses (250-10,000 cells) in 100 µl of cold (4°C) FCB and kept on ice. High-concentration (HC) Matrigel matrix (Corning, 354262), was thawed on ice, diluted (1:2) with ice cold FCB, and finally added at 1:1 ratio to the suspensions of sorted cells (100 µl of diluted HC Matrigel + 100 µL of sorted cells in FCB) for a total volume of 200 µl/injection aliquot. Each aliquot of sorted cells admixed with HC Matrigel (200 µl) was then injected subcutaneously (s.c.) in an NSG mice using 23 G x 1 ¼ needles. Paired sets of autologous CD49f^high^/KIT^neg^ and CD49f^low^/KIT^+^ cell populations were injected in parallel, in the s.c. tissue of the left and right flanks of the same animals, in order to exclude confounding effects from possible individual variabilities in the animals’ immune-competence. Animals were assessed weekly for the presence or absence of tumors, and upon tumor formation, tumor volume was measured weekly using the following formula:

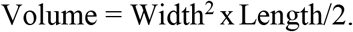

Animals were euthanized when tumors reached a maximum diameter of 2.0 cm, or followed for a minimum of 10 months, to exclude tumor engraftment. Upon euthanasia, animals were dissected and the s.c. tissues of both flanks was examined to exclude the presence of sub-palpable tumors. *Extreme Limiting Dilution Analysis* (ELDA) of tumorigenicity data was performed using a public online platform provided by the *Walter and Eliza Hall Institute* (WEHI) (http://bioinf.wehi.edu.au/software/elda)^29^. Tumors originated from the injection of purified preparations of either CD49f^high^/KIT^neg^ or CD49f^low^/KIT^+^ cells were analyzed by flow-cytometry and IHC, to evaluate their cell composition. Differences in the ratio of CD49f^high^/KIT^neg^ cells and CD49f^low^/KIT^+^ cells observed between parent tumors and tumors generated from purified populations were tested for statistical significance using an ordinary one-way ANOVA with Dunnett’s multiple comparisons test.

### *Three-dimensional* (3D) organoid cultures

One day prior to organoid plating, irradiated (100 Gy) feeder cells, consisting of a 1:1 mixture of L-Wnt-3A mouse fibroblasts (ATCC, CRL-2647), and R-Spondin1-HEK-293T cells (Trevigen, 3710-001-K), were thawed and plated at a density of 400,000 cells/well in a 24-well plate, after resuspension in a “feeder medium”, consisting of DMEM (Corning, 10-013-CV) containing 10% FBS (VWR, 89510-194), 100 U/mL penicillin and 100 µg/mL streptomycin (Millipore Sigma, P4333), 2 mM L-alanyl-L-glutamine (Corning 25-015-CI), 1 mM sodium pyruvate (Gibco, 11360070), and 20 mM HEPES (Corning, 25-060-CI). After 24 hours, the feeder medium was replaced with an “organoid medium”, consisting of DMEM Nutrient Mixture F-12 HAM tissue-culture medium (Sigma, D8437), supplemented with 10% heat-inactivated FBS (VWR, 89510-194), 2 mM L-alanyl-L-glutamine (Corning 25-015-CI), 20 mM HEPES (Corning, 25-060-CI), 1 mM sodium pyruvate (Gibco, 11360070), 100 U/mL penicillin and 100 µg/mL streptomycin (Sigma, P4333), 1x Antibiotic Antimycotic Solution (Corning 30-004-Cl), 1x ITES media supplement (Lonza, 17-839Z), 10 mM Nicotinamide (Sigma, 72340), and 100 ug/ml Heparin (Millipore Sigma, H3393). On the day of organoid plating, ACC tissues were dissociated into small fragments, as described above. Tissue fragments were then serially filtered through a 100 µm and a 40 µm mesh strainer. After the second filtration step, fragments trapped by the 40 µm strainer (i.e. tissue fragments smaller than 100 µm, but larger than 40 µm), were gently washed from the mesh with cold disaggregation medium and pelleted by centrifugation (1500 RPM, 5 minutes). Fragments were then resuspended in “complete organoid medium”, i.e., organoid medium supplemented with 50 ng/ml hEGF (Stem Cell Technologies, 78006.2), 500 ng/ml hR-Spondin1 (R&D systems, 4645-RS), and 10 µM Y-27632 (R&D Systems, 1254), and plated in transwell inserts (Griener Thincert, 24 well, 0.4µM pore size, 662641) atop a polymerized layer (∼100 µL) of Matrigel (Corning 354234). Finally, transwells were placed in 24-well plates atop feeder cells and cultures were incubated at 37°C with 5% CO_2_.

### *In vitro* studies with direct and inverse agonist of *retinoic acid* (RA) signaling

Stock solutions of direct and inverse agonists of RA signaling, including *all-trans retinoic acid* (ATRA;Sigma R2625, 100 mM), Bexarotene (Tocris 5819, 100 mM), BMS493 (Tocris 3509, 10 mM), and AGN193109 (Sigma SML2034, 10 mM), were prepared in DMSO and stored at −20°C, protected from light. On the day of use, stock solutions were thawed and added to complete organoid medium, at appropriate concentrations (100 nM −10 µM). Due to the short half-life of retinoids, medium with retinoid compounds was kept for a maximum of 3 days at 4°C and changed daily for the duration of culture (7 days).

### Analysis of organoid cell composition (histology, FACS, brightfield microscopy)

Organoid cultures established from human ACCs were dissociated from Matrigel by incubation with a solution of 2 mg/mL Dispase-II (Thermo Fisher, 17105041) and 200 U/mL Collagenase-III (Worthington, LS004183) in DPBS at 37°C for 15 minutes. Organoid suspensions were then transferred to 1.5 mL plastic tubes and pelleted by centrifugation (10,000 RPM, 2 minutes). Excess Matrigel and disaggregation solution were carefully aspirated. To dissolve remaining Matrigel, organoid pellets were briefly (3 minutes) resuspended in 0.25% Trypsin at 37°C, followed by a wash step with cell culture medium containing 10% FBS. To prepare organoids for immunohistochemistry, organoids were pelleted by centrifugation and fixed in 10% formalin for 4-12 hours. Fixed organoids were embedded in paraffin blocks, from which 4 µm tissue-sections were cut and stained following protocols identical to those used for tumor tissues (described above). To prepare organoids for FACS, organoid pellets were resuspended in disaggregation medium containing DNase-I (100 U/mL), Collagenase-III (200 U/ml), and Hyaluronidase (100 U/ml) and incubated at 37 °C for 20-30 minutes. Disaggregated organoids were then pelleted and incubated in 0.25% Trypsin for 10-15 minutes to generate a single cell suspension. Dissociated cells were washed with cell culture medium containing 10% FBS to inhibit Trypsin activity, followed by blocking with human IgGs (5mg/ml) and staining with antibodies. Differences in the percentage of CD49f^high^/KIT^neg^ and CD49f^low^/KIT^+^ cells in ACC organoids treated with retinoid compounds were tested for statistical significance using a one-way ANOVA with Dunnett’s multiple comparisons test. Brightfield images of organoid cultures were acquired using a Cytation 5 Cell Imaging Reader (BioTek) at 4x magnification. Images of *hematoxylin and eosin* (H&E) or IHC-stained organoids were acquired using a Nikon Eclipse E600 microscope with NIS-Elements Software (version 5.21). Brightness and contrast were adjusted uniformly throughout whole images using Adobe Photoshop (22.5.0 Release).

### Two-dimensional (2D) *in vitro* culture of purified preparations of myoepithelial-like and ductal-like cells

CD49f^high^/KIT^neg^ and CD49f^low^/KIT^+^ cell populations were sorted by FACS from ACC xenografts (ACCX5M1). Sorted cell populations were resuspended in 100 µl of complete organoid medium supplemented with either DMSO, ATRA (10 µM) or BMS493 (10 µM). Cells were plated in 96-well plates (30,000 cells/well) and medium was changed daily for the duration of treatment (1 week).

### Cell viability assays

2D cultures of ACCX5M1 cells and 3D organoid cultures of ACCX9 and ACCX11 were grown in 96-well black plates with optically clear bottoms (Thermo Scientific; 165305), and then treated with either retinoids (ATRA 10 µM, BMS493 10 µM) or DMSO alone (1:1000) for one week. On the final day of treatment, a 20% solution of alamarBlue HS cell viability reagent (Invitrogen A50100) was prepared in complete organoid medium and added to each well (final concentration = 10% alamarBlue reagent). The samples were incubated overnight (18-24 hours), at 37°C and protected from light^61^. Sample fluorescence was measured using a Cytation 5 Cell Imaging Reader (BioTek; ex/em 530/590). AlamarBlue fluorescence values were normalized to the mean of DMSO-treated control samples and differences in relative viability were tested for statistical significance using a two-tailed Student’s t-test.

### Evaluation of the *in vivo* anti-tumor activity of BMS493

BMS493 (Tocris, 3509) was resuspended in DMSO (stock concentration: 50 mg/mL) and stored at −20°C in single-use aliquots (20 µl). On the day of *in vivo* administration, single-use aliquots were thawed, and BMS493 was further diluted to a concentration of 2 mg/mL in DPBS supplemented with 0.15M *hydroxypropyl β-cyclodextrin* (HP-β-CD; Cayman Chemicals 16169), for a total volume of 0.5 mL per dose (1 mg/dose). To facilitate compound dissolution, diluted BMS493 or DMSO was warmed at 37°C for 10 minutes prior to injection. Mice were treated with either BMS493 or vehicle alone (DMSO, 0.15 M HP-β-CD) by intraperitoneal injection, according to two treatment regimens: Regimen 1: 3 doses/week (Treat M, W, F) for three weeks (9 mg total dose); Regimen 2: 4 doses/week (Treat M, Tu, Th, F), for three weeks (12 mg total dose). Tumor volume was measured weekly, mice were weighted twice per week, and animals were monitored daily. Tumor volume was calculated using the following formula:

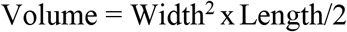

To enable robust comparisons across different treatment groups, tumor volumes were normalized to their starting values, and reported as fold-increases over time. Tumor growth rates were then calculated assuming exponential kinetics, as described by Hather et al^62^. Differences in normalized tumor volumes at individual time points were tested for statistical significance using a two-tailed Student’s t-test, while differences between growth rates (i.e., Log10(fold-increase in tumor volume)/Time(days)) were tested for statistical significance using a two-tailed Welch’s t-test (i.e., assuming unequal variance).

## AUTHOR CONTRIBUTIONS (CRediT)

**Sara Viragova:** Conceptualization, Investigation, Formal analysis, Methodology, Validation, Visualization, Writing-original draft

**Luis Aparicio:** Conceptualization, Investigation, Formal analysis, Methodology, Data Curation, Visualization, Writing- review & editing

**Junfei Zhao:** Formal analysis, Software, Data curation

**Luis E. Valencia Salazar**: Investigation, Validation

**Alexandra Schurer:** Investigation, Validation

**Anika Dhuri:** Investigation, Validation

**Debashis Sahoo:** Formal analysis, Software, Data curation

**Christopher Moskaluk:** Methodology, Resources

**Raul Rabadan:** Funding acquisition, Supervision, Resources

**Piero Dalerba:** Conceptualization, Resources, Supervision, Funding acquisition, Validation, Investigation, Visualization, Project administration, Writing - original draft, Writing- review & editing

## EXTENDED DATA FIGURES

**Extended Data Figure 1:**
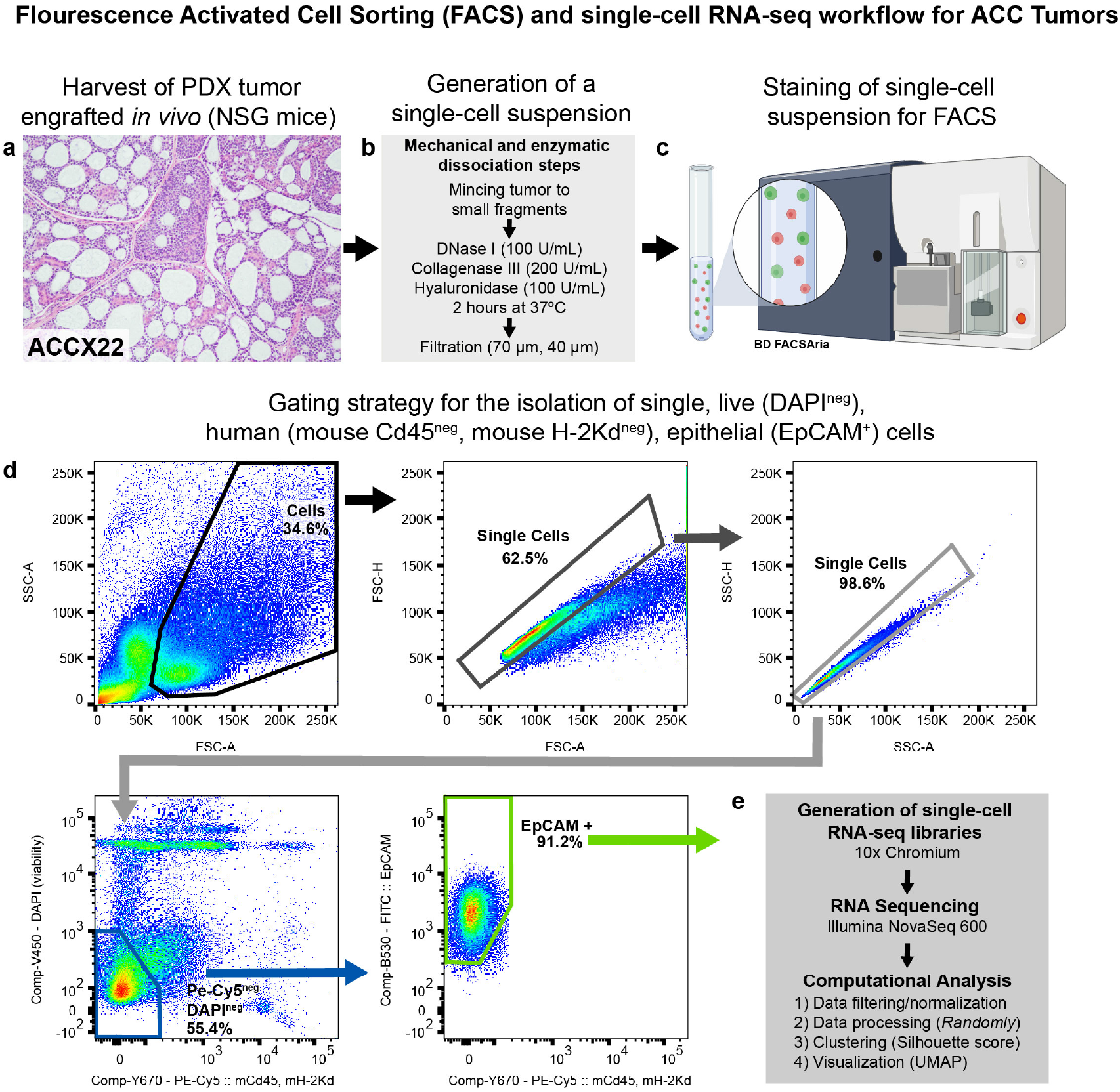
Workflow of *single-cell RNA-sequencing* (scRNA-seq) experiment to analyze the cell composition of *patient-derived xenograft* (PDX) line ACCX22. **(a)** ACCX22 tumor was first stained with *hematoxylin and eosin* (H&E) and confirmed to retain a cribriform histology, characteristic of well-differentiated (Grade 1) ACCs. **(b)** Tumor tissues were cut into small fragments with scissors, minced, and dissociated into a single-cell suspension by enzymatic digestion (DNase I, Collagenase III, Hyaluronidase). **(c)** Single-cell suspensions were analyzed and sorted using the BD FACSAria. **(d)** Gating strategy to isolate single, live (DAPI^neg^), human (mouse Cd45^neg^, mouse H-2Kd^neg^), epithelial (EpCAM^+^) cells from ACCX22 PDX model for single cell analysis. **(e)** Overview of experimental and analysis steps for scRNA-seq. Single-cell libraries were prepared using the 10x Chromium system (Single Cell 3’ v3 chemistry), and sequenced using the NovaSeq 6000 platform (Illumina). Single-cell data was filtered to retain cells with >500 genes/cell and normalized (log2(1+TPM)). Gene-expression data was processed to remove signals attributable to stochastic variation (noise) using the *Randomly* package, then used to sub-group cells into distinct clusters, which were visualized using *Uniform Manifold Approximation and Projection* (UMAP). Schematic in panel (c) was created with BioRender.com

**Extended Data Figure 2:**
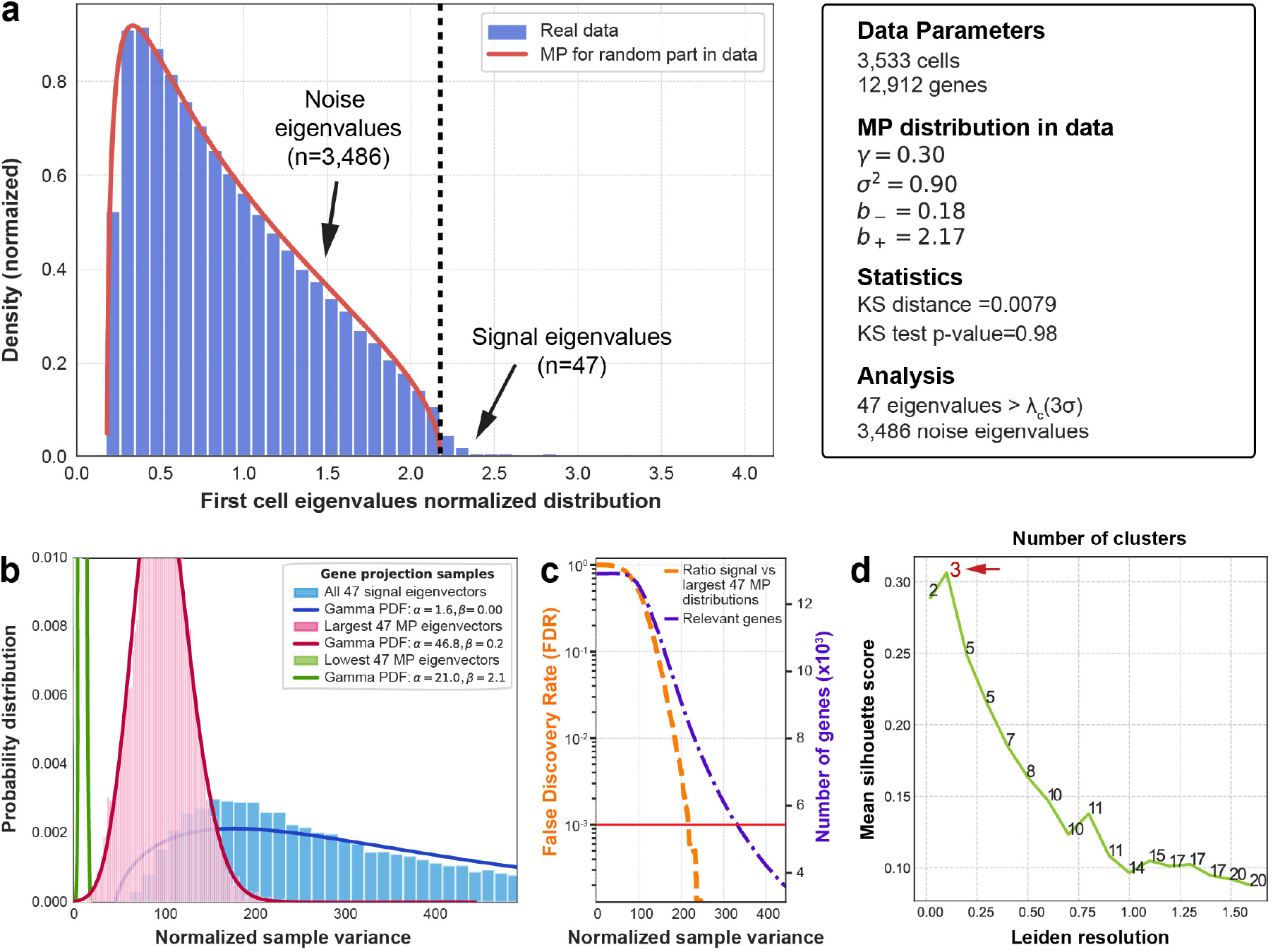
Computational analysis of *single-cell RNA-sequencing* (scRNA-seq) data obtained from *patient-derived xenograft* (PDX) line ACCX22. **(a)** Spectral distribution (blue histogram) of the Wishart matrix for 3,533 cells identified as sequenced at sufficient depth (>500 expressed genes), after elimination of sparsity-induced signal. After fitting the *Marchenko-Pastur* (MP) distribution to the data (red curve), 47 eigenvalues (1.3%; n=47/3,533) are identified as lying outside the MP distribution, and thus as corresponding to eigenvectors that carry informative signal. **(b)** Study of the chi-squared test for the variance (normalized sample variance) of each gene’s projection into noise and signal eigenvectors. The blue distribution was generated based on the 47 signal-like eigenvectors, the pink distribution based on the eigenvectors corresponding to the highest 47 eigenvalues within the MP distribution, and the green distribution based on the eigenvectors corresponding to the smallest 47 MP eigenvalues within the MP distribution. **(c)** Distribution of the number of genes identified as mostly responsible for the signal (purple line) and of their *false discovery rate* (FDR; orange line) as a function of the normalized sample variance. The FDR is calculated as the ratio of the blue and pink distributions in panel b. Approximately, 5,500 genes are found responsible for the signal when adopting an FDR threshold of ≤0.001 (horizontal red line). **(d)** Relationship between mean *silhouette score*, number of candidate cell clusters and level of resolution imposed through the Leiden clustering algorithm (Wolf et al., *Genome Biology*, 19:1-5, 2018), after removal of signals attributable to noise using *Randomly*. The optimal clustering solution (i.e., the clustering solution with the highest mean silhouette score) corresponds to three clusters. A complete description of this computational pipeline was previously published (Aparicio et al., *Patterns*, 1:100035, 2020).

**Extended Data Figure 3:**
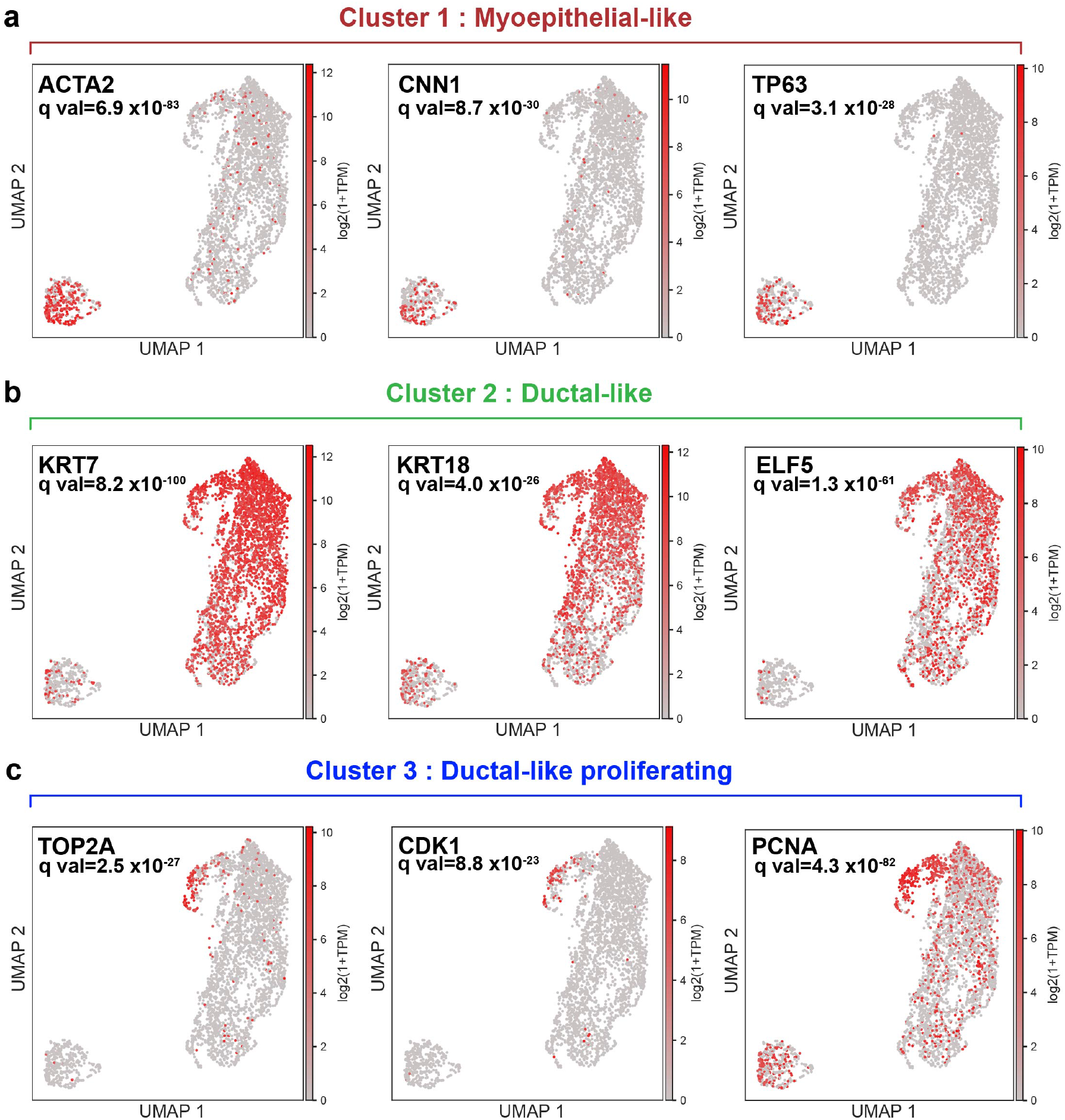
Distribution of expression levels for myoepithelial-specific, ductal-specific and proliferation-specific biomarkers in scRNA-seq data from *patient-derived xenograft* (PDX) line ACCX22. **(a)** Expression of genes encoding for myoepithelial markers, including *smooth muscle actin alpha 2* (ACTA2), *calponin* (CNN1), and *tumor protein p63* (TP63) is enriched in cells belonging to Cluster 1. **(b)** Expression of genes encoding for ductal/luminal markers, including *keratin 7* (KRT7), *keratin 18* (KRT18), and *E74-like ETS transcription factor 5* (ELF5) is enriched in cells belonging to Clusters 2 and 3. **(c)** Expression of genes encoding for proliferation markers, including *DNA topoisomerase II alpha* (TOP2A), *cyclin-dependent kinase 1* (CDK1), and *proliferating cell nuclear antigen* (PCNA) is enriched in cells belonging to Cluster 3. Q values represent the result of a t-test with a Benjamini-Hochberg multiple hypothesis testing correction, comparing the mean expression level of the marker of interest within the cluster that preferentially expresses it vs. mean expression level of the marker in all other cells.

**Extended Data Figure 4:**
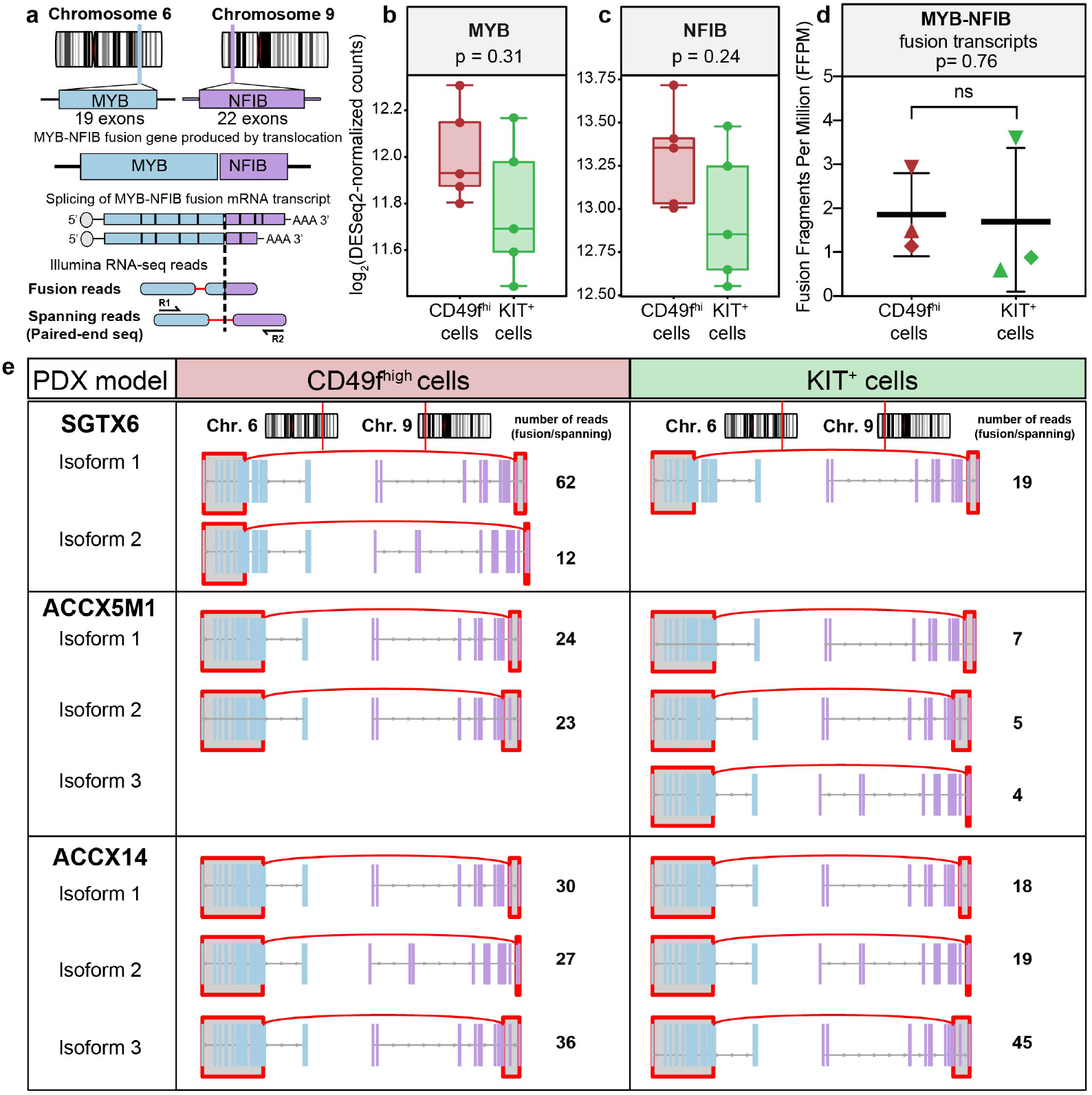
Analysis of *MYB-NFIB* fusion transcripts in myoepithelial-like (CD49f^high^/KIT^neg^) and ductal-like (CD49f^low^/KIT^+^) cells. **(a)** Schematic description of *MYB-NFIB* fusion genes and their chimeric mRNA transcripts. *MYB-NFIB* mRNAs undergo alternative splicing, predominantly in the *NFIB* portion of the fusion gene, thus generating distinct chimeric isoforms. In RNA-seq data, fusion transcripts can be identified as chimeric reads (i.e., reads encompassing two distinct genes) or, in paired-end sequencing, as spanning reads (i.e., paired reads mapping to different genes), using the STAR-fusion software. **(b-c)** Analysis of *MYB* and *NFIB* gene expression levels in RNA-seq data from five bi-phenotypic PDX models (ACCX5M1, ACCX6, ACCX14, ACCX22, SGTX6). Both genes were detected at high levels in both CD49f^high^/KIT^neg^ and CD49f^low^/KIT^+^ cell populations. Differences in expression levels were not statistically significant (Student’s t-test for paired samples, two-tailed). **(d)** *MYB-NFIB* fusion fragments per million (FFPM) transcripts were detected in both CD49f^high^/KIT^neg^ and CD49f^low^/KIT^+^ cells from three tumors with chromosomal translocations yielding chimeric mRNAs (ACCX5M1, ACCX14, SGTX6). Differences in expression levels of fusion transcripts were not statistically significant (Student’s t-test for paired samples, two-tailed). **(e)** In tumors with chromosomal translocations yielding chimeric mRNAs (ACCX5M1, ACCX14, SGTX6), the repertoire of *MYB-NFIB* splicing isoforms appeared similar in CD49f^high^/KIT^neg^ and CD49f^low^/KIT^+^ cells. Autologous pairs shared the same *MYB* breakpoint of parent tumors, thus confirming their shared origin from a common cellular ancestor.

**Extended Data Figure 5:**
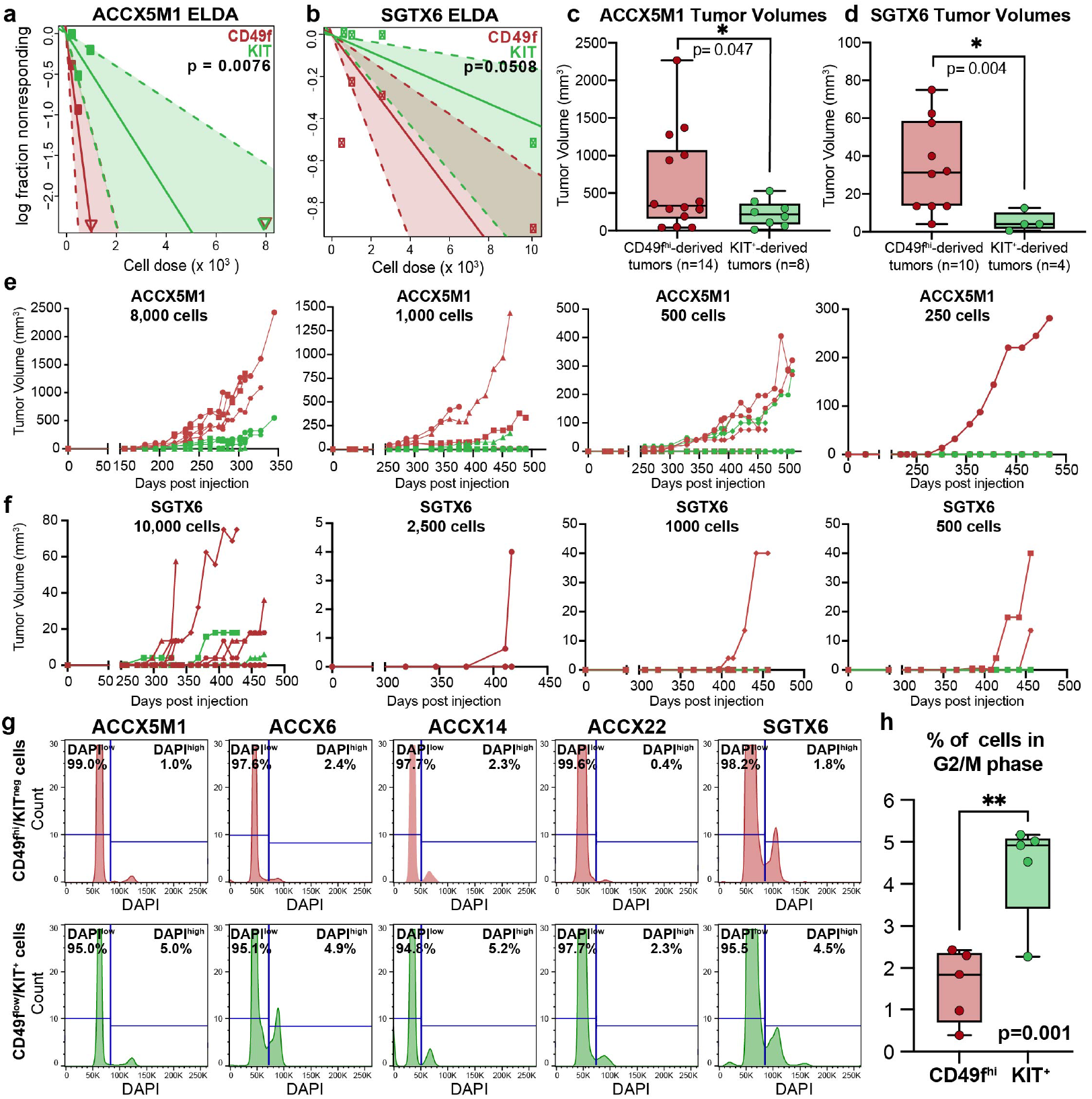
Comparison of tumorigenic capacity and cell cycle distribution of CD49f^high^/KIT^neg^ and CD49f^low^/KIT^+^ cells from human ACCs. **(a-b)** *Extreme Limiting Dilution Analysis* (ELDA) of tumorigenicity data from CD49f^high^/KIT^neg^ and CD49f^low^/KIT^+^ cells isolated by flow cytometry from the ACCX5M1 (a) and SGTX6 (b) PDX lines, reported using a Log-fraction plot. The slope of fitted lines (red: CD49f^high^/KIT^neg^; green: CD49f^low^/KIT^+^) represents the log-active cell fraction (shaded areas: 95% confidence interval). **(c-d)** Comparison of tumor volumes measured at euthanasia in animals engrafted with CD49f^high^/KIT^neg^ (red) and CD49f^low^/KIT^+^ (green) cells purified by flow cytometry from the ACCX5M1 (c) and SGTX6 (d) PDX lines. Tumors originated from CD49f^high^/KIT^neg^ cells reached higher volumes than tumors originated from CD49f^low^/KIT^+^ cells (Welch’s t-test, two-tailed). **(e-f)** Growth curves of individual tumors originated from *in vivo* injection of CD49f^high^/KIT^neg^ (red) and CD49f^low^/KIT^+^ (green) cells, isolated by flow cytometry from the ACCX5M1 (e) and SGTX6 (f) PDX lines. Tumors appeared between 150-300 days (ACCX5M1) and 250-450 days (SGTX6) post-engraftment. **(g)** Analysis of cell cycle distribution of CD49f^high^/KIT^neg^ and CD49f^low^/KIT^+^ cells from five bi-phenotypic ACC PDX lines. **(h)** The percentage of cells in the G2/M phase of the cell cycle (DAPI^high^) was lower in CD49f^high^/KIT^neg^ as compared to CD49f^low^/KIT^+^ cells (p=0.001; Student t-test for paired samples, two-tailed).

**Extended Data Figure 6:**
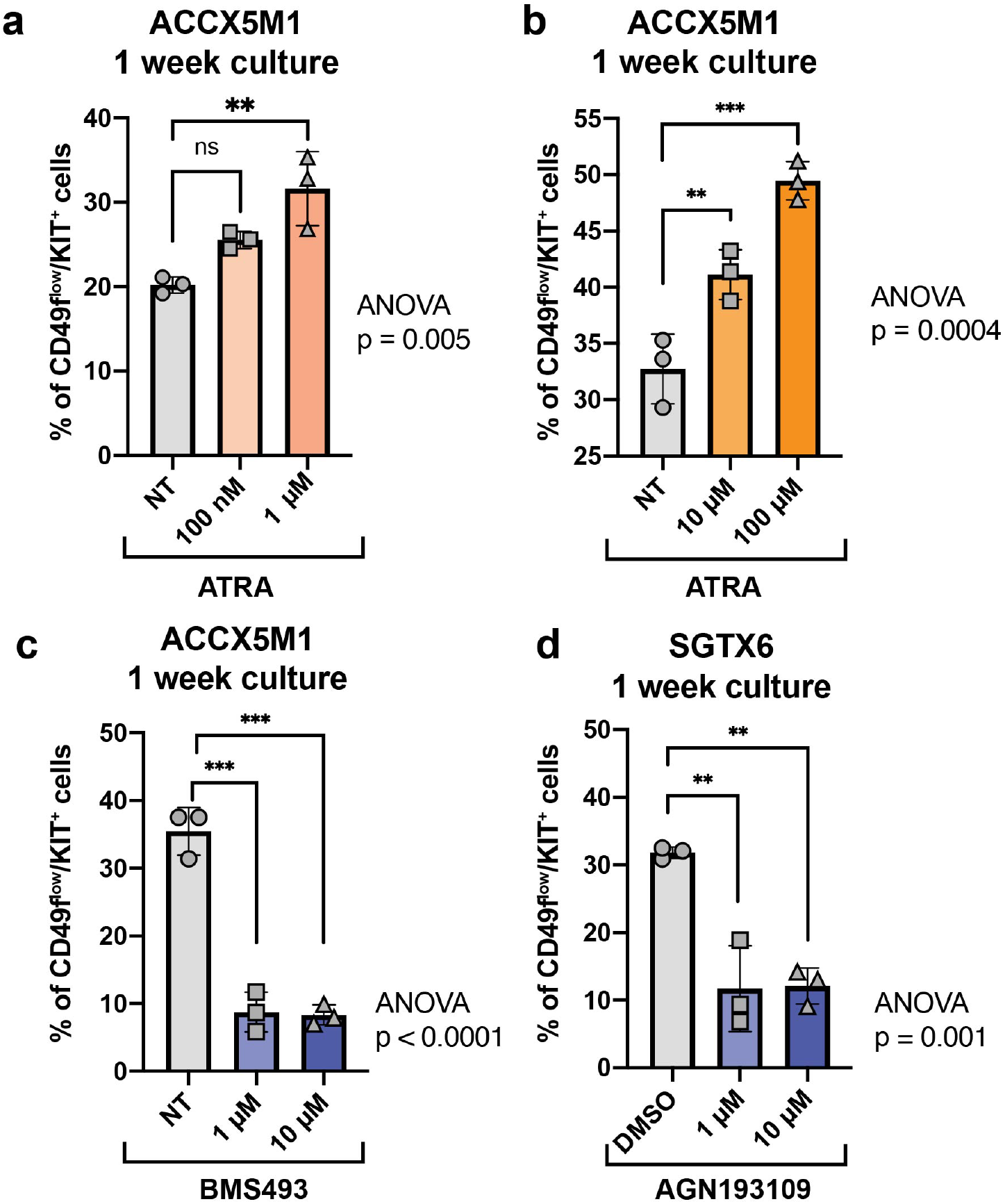
Dose-response effects of pharmacological modulators of *retinoic acid* (RA) signaling on the percentage of CD49f^low^/KIT^+^ cells in organoids established from PDX lines representative of bi-phenotypic ACCs. **(a-b)** Treatment with *all-trans retinoic acid* (ATRA), an agonist of RA signaling, induces a dose-dependent increase in the percentage of CD49f^low^/KIT^+^ cells in 3D organoid cultures established from the ACCX5M1 PDX line. The increase in the percentage of CD49f^low^/KIT^+^ cells is already observed at the 0.1-1 µM concentration range (a), and becomes progressively more prominent as the concentration is raised to the 10-100 µM concentration range (b). **(c-d)** Treatment with either BMS493 or AGN19310, two inverse agonists (i.e., inhibitors) of RA signaling, results in a profound reduction in the percentage of CD49f^low^/KIT^+^ cells in 3D organoid cultures established from either the ACCX5M1 (c; BMS43) or SGTX6 (d; AGN193109) PDX lines, even when administered at low doses (1 µM). Organoid cultures were treated for one week and analyzed by flow cytometry (n=3 wells/condition). Changes in the percentage of CD49f^low^/KIT^+^ cells were tested for statistical significance using one-way ANOVA with Dunnett’s multiple comparisons test against untreated (NT) or DMSO-treated controls (ns = non-significant, *p<0.05, **p<0.01, ***p<0.001).

**Extended Data Figure 7:**
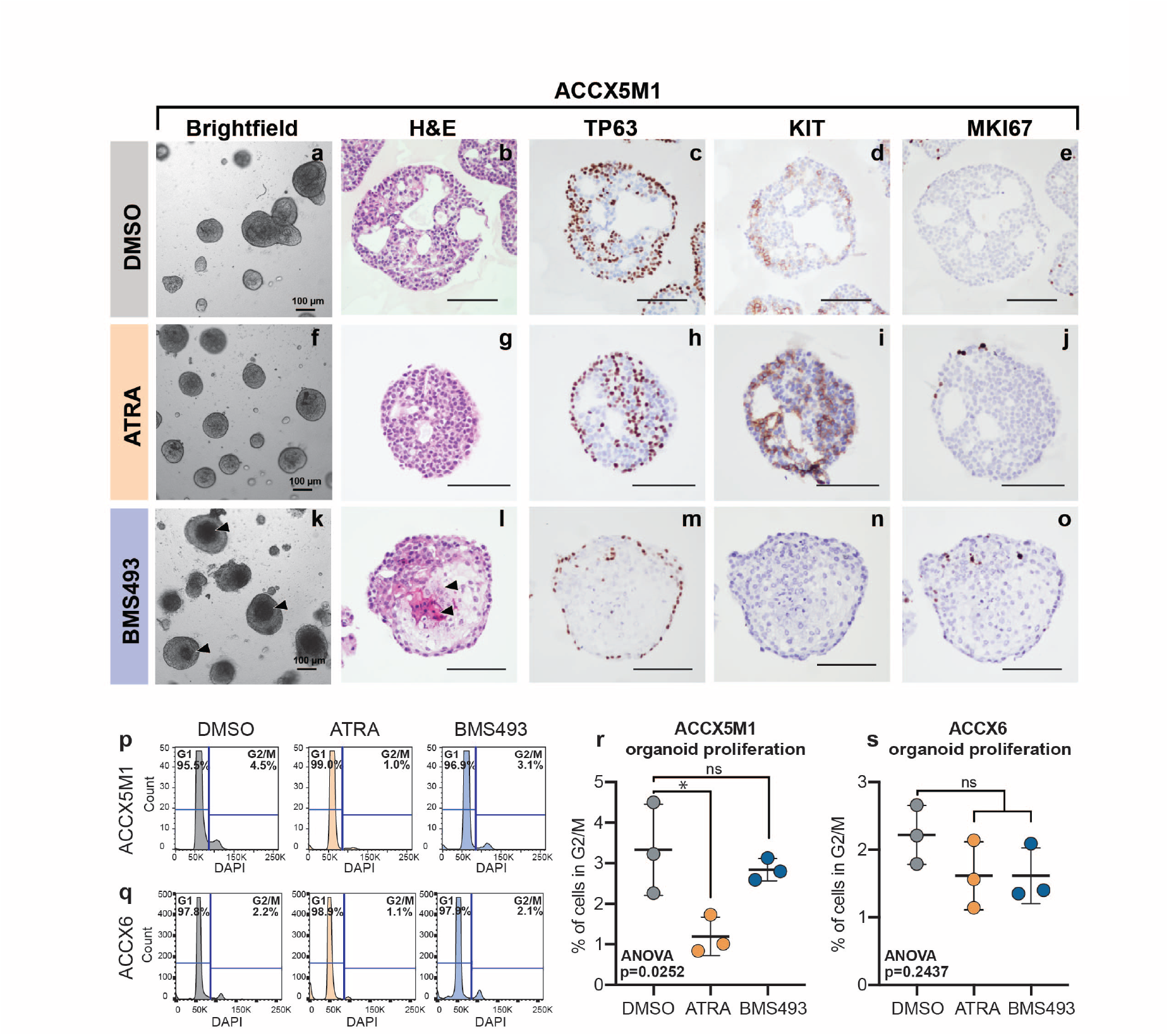
Perturbation of *retinoic acid* (RA) signaling does not induce proliferation in ACC organoids. **(a-o)** Analysis of organoid morphology and histology, following treatment with either activators (ATRA; direct agonist) or inhibitors (BMS493; inverse agonist) of RA signaling. Organoids were established from a PDX line with bi-phenotypic histology (ACCX5M1) and treated for one week with either DMSO (a-e), 10 µM ATRA (f-j) or 10 µM BMS493 (k-o). Scale bars = 100 µm. Treatment with ATRA did not change organoid morphology (f), but increased the number of KIT^+^ cells (i), as visualized by *immunohistochemistry* (IHC). Treatment with BMS493 caused a dramatic change in organoid morphology, characterized by the appearance of dense areas in the organoid centers, when observed using bright-field microscopy (k, arrowheads). When organoids were stained with hematoxylin and eosin (H&E), these areas consisted of an eosin-rich material, with apoptotic nuclei (l, arrowheads). **(p-q)** Analysis by flow cytometry of cell cycle distribution in organoids established from ACCX5M1 (p) and ACCX6 (q) PDX lines, following 1 week of treatment with either DMSO, ATRA (10 µM) or BMS493 (10µM). **(r-s)** Treatment with either ATRA or BMS493 did not increase the percentage of cells in the G2/M phase of the cell-cycle in ACCX5M1 (r) and ACCX6 (s) organoids. Experiments included at least three replicates (n=3 wells/condition). Error bars: mean +/- standard deviation. Differences in the percentage of cells in the G2/M phase were tested for statistical significance using one-way ANOVA with Dunnett’s multiple comparisons test (ns=non-significant, *p<0.05).

**Extended Data Figure 8:**
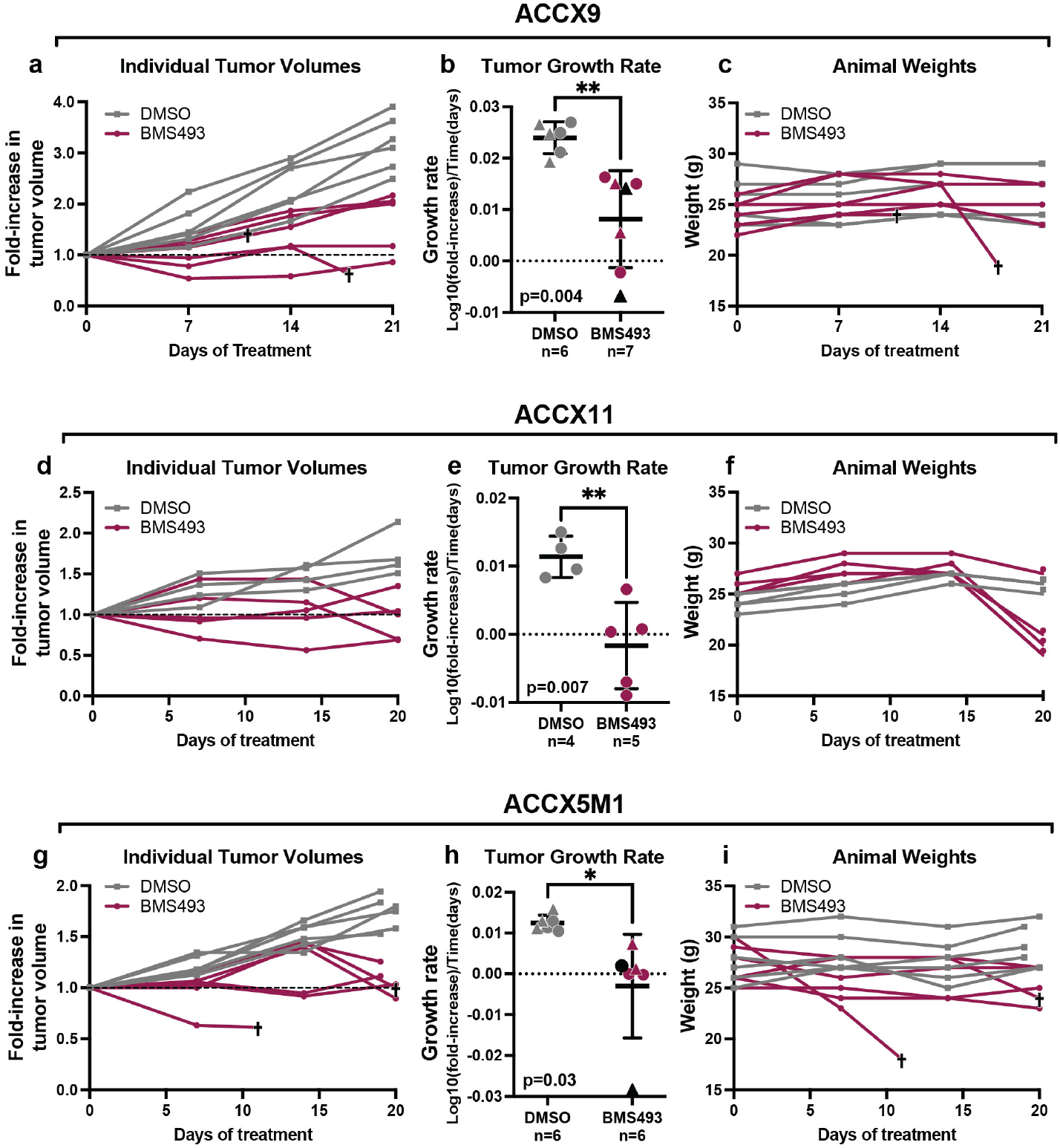
*In vivo* anti-tumor activity and toxicity of BMS493. **(a)** Individual growth curves of ACCX9 tumors treated with DMSO (gray) or BMS493 (magenta). Cross symbols (†) denote two animals who were sacrificed early, due to deterioration of their health condition. **(b)** Growth rates of ACCX9 tumors treated with either DMSO (n=6) or BMS493 (n=7). Two treatment cohorts are identified by different symbols (circles = cohort 1; triangles = cohort 2). Differences in mean growth rates were statistically significant (Welch’s t-test; **p<0.01). Black symbols denote the two animals who were sacrificed because of a deterioration of their health condition. **(c)** Individual animal weights over the course of *in vivo* treatment with BMS493 (†: animals sacrificed due to deterioration of health conditions). **(d)** Individual growth curves of ACCX11 tumors treated with DMSO (gray) or BMS493 (magenta). **(e)** Growth rates of ACCX11 tumors treated with either DMSO (n=4) or BMS493 (n=5). Differences in mean growth rates were statistically significant (Welch’s t-test; **p<0.01). **(f)** Individual animal weights over the course of *in vivo* treatment with BMS493. **(g)** Individual growth curves of ACCX5M1 tumors treated with DMSO (gray) or BMS493 (magenta). Cross symbols (†) denote two animals who either were sacrificed early due to deterioration of their general health or who died at the final time point. **(h)** Growth rates of ACCX5M1 tumors treated with either DMSO (n=6) or BMS493 (n=6). Two treatment cohorts are identified by different symbols (circles = cohort 1; triangles = cohort 2). Differences in mean growth rates were statistically significant (Welch’s t-test; *p<0.05). Black symbols denote the two animals who either were sacrificed early due to deterioration of their general health or died at the final time point. **(i)** Individual animal weights over the course of treatment (†: animals sacrificed or found dead).

## REFERENCES

1. Moskaluk, C.A. Adenoid cystic carcinoma: clinical and molecular features. Head and neck pathology 7, 17–22 (2013).

2. Gao, M., Hao, Y., Huang, M., Ma, D., Luo, H., Gao, Y., Peng, X. & Yu, G. Clinicopathological study of distant metastases of salivary adenoid cystic carcinoma. International journal of oral and maxillofacial surgery 42, 923–928 (2013).

3. Di Villeneuve, L., Souza, I.L., Tolentino, F.D.S., Ferrarotto, R. & Schvartsman, G. Salivary Gland Carcinoma: Novel Targets to Overcome Treatment Resistance in Advanced Disease. Frontiers in Oncology 10, 2097 (2020).

4. Laurie, S.A., Ho, A.L., Fury, M.G., Sherman, E. & Pfister, D.G. Systemic therapy in the management of metastatic or locally recurrent adenoid cystic carcinoma of the salivary glands: a systematic review. The lancet oncology 12, 815–824 (2011).

5. Ho, A.S., Kannan, K., Roy, D.M., Morris, L.G., Ganly, I., Katabi, N., Ramaswami, D., Walsh, L.A., Eng, S. & Huse, J.T. The mutational landscape of adenoid cystic carcinoma. Nature genetics 45, 791–798 (2013).

6. Stephens, P.J., Davies, H.R., Mitani, Y., Van Loo, P., Shlien, A., Tarpey, P.S., Papaemmanuil, E., Cheverton, A., Bignell, G.R., Butler, A.P., Gamble, J., Gamble, S., Hardy, C., Hinton, J., Jia, M., Jayakumar, A., Jones, D., Latimer, C., McLaren, S., McBride, D.J., Menzies, A., Mudie, L., Maddison, M., Raine, K., Nik-Zainal, S., O’Meara, S., Teague, J.W., Varela, I., Wedge, D.C., Whitmore, I., Lippman, S.M., McDermott, U., Stratton, M.R., Campbell, P.J., El-Naggar, A.K. & Futreal, P.A. Whole exome sequencing of adenoid cystic carcinoma. J Clin Invest 123, 2965–2968 (2013).

7. Azumi, N. & Battifora, H. The cellular composition of adenoid cystic carcinoma. An immunohistochemical study. Cancer 60, 1589–1598 (1987).

8. Li, N., Xu, L., Zhao, H., El-Naggar, A.K. & Sturgis, E.M. A comparison of the demographics, clinical features, and survival of patients with adenoid cystic carcinoma of major and minor salivary glands versus less common sites within the Surveillance, Epidemiology, and End Results registry. Cancer 118, 3945–3953 (2012).

9. Du, F., Zhou, C.-X. & Gao, Y. Myoepithelial differentiation in cribriform, tubular and solid pattern of adenoid cystic carcinoma: A potential involvement in histological grading and prognosis. Annals of Diagnostic Pathology 22, 12–17 (2016).

10. Dalerba, P., Cho, R.W. & Clarke, M.F. Cancer stem cells: models and concepts. Annu. Rev. Med. 58, 267–284 (2007).

11. Dalerba, P., Kalisky, T., Sahoo, D., Rajendran, P.S., Rothenberg, M.E., Leyrat, A.A., Sim, S., Okamoto, J., Johnston, D.M. & Qian, D. Single-cell dissection of transcriptional heterogeneity in human colon tumors. Nature biotechnology 29, 1120 (2011).

12. Aparicio, L., Bordyuh, M., Blumberg, A.J. & Rabadan, R. A random matrix theory approach to denoise single-cell data. Patterns 1, 100035 (2020).

13. Moskaluk, C.A., Baras, A.S., Mancuso, S.A., Fan, H., Davidson, R.J., Dirks, D.C., Golden, W.L. & Frierson, H.F. Development and characterization of xenograft model systems for adenoid cystic carcinoma. Lab Invest 91, 1480–1490 (2011).

14. Dalerba, P., Dylla, S.J., Park, I.K., Liu, R., Wang, X., Cho, R.W., Hoey, T., Gurney, A., Huang, E.H., Simeone, D.M., Shelton, A.A., Parmiani, G., Castelli, C. & Clarke, M.F. Phenotypic characterization of human colorectal cancer stem cells. Proc Natl Acad Sci U S A 104, 10158–10163 (2007).

15. Love, M.I., Huber, W. & Anders, S. Moderated estimation of fold change and dispersion for RNA-seq data with DESeq2. Genome biology 15, 550 (2014).

16. Prasad, M.L., Barbacioru, C.C., Rawal, Y.B., Husein, O. & Wen, P. Hierarchical cluster analysis of myoepithelial/basal cell markers in adenoid cystic carcinoma and polymorphous low-grade adenocarcinoma. Modern Pathology 21, 105–114 (2008).

17. Wu, H.-M., Ren, G.-X., Wang, L.-Z., Zhang, C.-Y., Chen, W.-T. & Guo, W. Expression of podoplanin in salivary gland adenoid cystic carcinoma and its association with distant metastasis and clinical outcomes. Molecular medicine reports 6, 271–274 (2012).

18. Oakes, S.R., Naylor, M.J., Asselin-Labat, M.-L., Blazek, K.D., Gardiner-Garden, M., Hilton, H.N., Kazlauskas, M., Pritchard, M.A., Chodosh, L.A., Pfeffer, P.L., Lindeman, G.J., Visvader, J.E. & Ormandy, C.J. The Ets transcription factor Elf5 specifies mammary alveolar cell fate. Genes & Development 22, 581–586 (2008).

19. Iglesias, J.M., Cairney, C.J., Ferrier, R.K., McDonald, L., Soady, K., Kendrick, H., Pringle, M.-A., Morgan, R.O., Martin, F. & Smalley, M.J. Annexin A8 identifies a subpopulation of transiently quiescent c-kit positive luminal progenitor cells of the ductal mammary epithelium. PloS one 10, e0119718 (2015).

20. Lim, E., Vaillant, F., Wu, D., Forrest, N.C., Pal, B., Hart, A.H., Asselin-Labat, M.L., Gyorki, D.E., Ward, T., Partanen, A., Feleppa, F., Huschtscha, L.I., Thorne, H.J., kConFab, Fox, S.B., Yan, M., French, J.D., Brown, M.A., Smyth, G.K., Visvader, J.E. & Lindeman, G.J. Aberrant luminal progenitors as the candidate target population for basal tumor development in BRCA1 mutation carriers. Nat Med 15, 907–913 (2009).

21. Nguyen, Q.H., Pervolarakis, N., Blake, K., Ma, D., Davis, R.T., James, N., Phung, A.T., Willey, E., Kumar, R. & Jabart, E. Profiling human breast epithelial cells using single cell RNA sequencing identifies cell diversity. Nature communications 9, 1–12 (2018).

22. Haas, B.J., Dobin, A., Li, B., Stransky, N., Pochet, N. & Regev, A. Accuracy assessment of fusion transcript detection via read-mapping and de novo fusion transcript assembly-based methods. Genome biology 20, 1–16 (2019).

23. Persson, M., Andrén, Y., Mark, J., Horlings, H.M., Persson, F. & Stenman, G. Recurrent fusion of MYB and NFIB transcription factor genes in carcinomas of the breast and head and neck. Proceedings of the National Academy of Sciences 106, 18740–18744 (2009).

24. Mitani, Y., Li, J., Rao, P.H., Zhao, Y.-J., Bell, D., Lippman, S.M., Weber, R.S., Caulin, C. & El-Naggar, A.K. Comprehensive analysis of the MYB-NFIB gene fusion in salivary adenoid cystic carcinoma: Incidence, variability, and clinicopathologic significance. Clinical cancer research 16, 4722–4731 (2010).

25. West, R.B., Kong, C., Clarke, N., Gilks, T., Lipsick, J., Cao, H., Kwok, S., Montgomery, K.D., Varma, S. & Le, Q.-T. MYB expression and translocation in adenoid cystic carcinomas and other salivary gland tumors with clinicopathologic correlation. The American journal of surgical pathology 35, 92 (2011).

26. Brill, L.B., Kanner, W.A., Fehr, A., Andrén, Y., Moskaluk, C.A., Löning, T., Stenman, G. & Frierson, H.F. Analysis of MYB expression and MYB-NFIB gene fusions in adenoid cystic carcinoma and other salivary neoplasms. Modern pathology 24, 1169–1176 (2011).

27. Drier, Y., Cotton, M.J., Williamson, K.E., Gillespie, S.M., Ryan, R.J.H., Kluk, M.J., Carey, C.D., Rodig, S.J., Sholl, L.M., Afrogheh, A.H., Faquin, W.C., Queimado, L., Qi, J., Wick, M.J., El-Naggar, A.K., Bradner, J.E., Moskaluk, C.A., Aster, J.C., Knoechel, B. & Bernstein, B.E. An oncogenic MYB feedback loop drives alternate cell fates in adenoid cystic carcinoma. Nat Genet 48, 265–272 (2016).

28. Panaccione, A., Zhang, Y., Ryan, M., Moskaluk, C.A., Anderson, K.S., Yarbrough, W.G. & Ivanov, S.V. MYB fusions and CD markers as tools for authentication and purification of cancer stem cells from salivary adenoid cystic carcinoma. Stem cell research 21, 160–166 (2017).

29. Hu, Y. & Smyth, G.K. ELDA: extreme limiting dilution analysis for comparing depleted and enriched populations in stem cell and other assays. Journal of immunological methods 347, 70–78 (2009).

30. Wright, D.M., Buenger, D.E., Abashev, T.M., Lindeman, R.P., Ding, J. & Sandell, L.L. Retinoic acid regulates embryonic development of mammalian submandibular salivary glands. Developmental Biology 407, 57–67 (2015).

31. Abashev, T.M., Metzler, M.A., Wright, D.M. & Sandell, L.L. Retinoic acid signaling regulates Krt5 and Krt14 independently of stem cell markers in submandibular salivary gland epithelium. Developmental Dynamics 246, 135–147 (2017).

32. Metzler, M.A., Raja, S., Elliott, K.H., Friedl, R.M., Tran, N.Q.H., Brugmann, S.A., Larsen, M. & Sandell, L.L. RDH10-mediated retinol metabolism and RARα-mediated retinoic acid signaling are required for submandibular salivary gland initiation. Development 145 (2018).

33. Mandelbaum, J., Shestopalov, I.A., Henderson, R.E., Chau, N.G., Knoechel, B., Wick, M.J. & Zon, L.I. Zebrafish blastomere screen identifies retinoic acid suppression of *MYB* in adenoid cystic carcinoma. The Journal of Experimental Medicine (2018).

34. Sun, B., Wang, Y., Sun, J., Zhang, C., Xia, R., Xu, S., Sun, S. & Li, J. Establishment of patient-derived xenograft models of adenoid cystic carcinoma to assess pre-clinical efficacy of combination therapy of a PI3K inhibitor and retinoic acid. American journal of cancer research 11, 773 (2021).

35. Dong, D., Ruuska, S.E., Levinthal, D.J. & Noy, N. Distinct roles for cellular retinoic acid-binding proteins I and II in regulating signaling by retinoic acid. Journal of Biological Chemistry 274, 23695–23698 (1999).

36. Schug, T.T., Berry, D.C., Shaw, N.S., Travis, S.N. & Noy, N. Dual transcriptional activities underlie opposing effects of retinoic acid on cell survival. Cell 129, 723 (2007).

37. Bastien, J. & Rochette-Egly, C. Nuclear retinoid receptors and the transcription of retinoid-target genes. Gene 328, 1–16 (2004).

38. Kam, R.K.T., Shi, W., Chan, S.O., Chen, Y., Xu, G., Bik-San Lau, C., Fung, K.P., Chan, W.Y. & Zhao, H. Dhrs3 protein attenuates retinoic acid signaling and is required for early embryonic patterning. Journal of Biological Chemistry 288, 31477–31487 (2013).

39. Du, C., Redner, R.L., Cooke, M.P. & Lavau, C. Overexpression of wild-type retinoic acid receptor alpha (RARalpha) recapitulates retinoic acid-sensitive transformation of primary myeloid progenitors by acute promyelocytic leukemia RARalpha-fusion genes. Blood 94, 793–802 (1999).

40. Sato, T., Stange, D.E., Ferrante, M., Vries, R.G., Van Es, J.H., Van Den Brink, S., Van Houdt, W.J., Pronk, A., Van Gorp, J. & Siersema, P.D. Long-term expansion of epithelial organoids from human colon, adenoma, adenocarcinoma, and Barrett’s epithelium. Gastroenterology 141, 1762–1772 (2011).

41. Shimono, Y., Mukohyama, J., Isobe, T., Johnston, D.M., Dalerba, P. & Suzuki, A. in Organoids 23–31 (Springer, 2016).

42. Panaccione, A., Chang, M.T., Carbone, B.E., Guo, Y., Moskaluk, C.A., Virk, R.K., Chiriboga, L., Prasad, M.L., Judson, B., Mehra, S., Yarbrough, W.G. & Ivanov, S.V. NOTCH1 and SOX10 are Essential for Proliferation and Radiation Resistance of Cancer Stem–Like Cells in Adenoid Cystic Carcinoma. Clinical Cancer Research 22, 2083 (2016).

43. Ferrarotto, R., Mitani, Y., McGrail, D.J., Li, K., Karpinets, T.V., Bell, D., Frank, S.J., Song, X., Kupferman, M.E. & Liu, B. Proteogenomic Analysis of Salivary Adenoid Cystic Carcinomas Defines Molecular Subtypes and Identifies Therapeutic Targets. Clinical Cancer Research (2020).

44. Ferrarotto, R., Mitani, Y., Diao, L., Guijarro, I., Wang, J., Zweidler-McKay, P., Bell, D., William Jr, W.N., Glisson, B.S. & Wick, M.J. Activating NOTCH1 mutations define a distinct subgroup of patients with adenoid cystic carcinoma who have poor prognosis, propensity to bone and liver metastasis, and potential responsiveness to Notch1 inhibitors. Journal of Clinical Oncology 35, 352 (2017).

45. Fordice, J., Kershaw, C., El-Naggar, A. & Goepfert, H. Adenoid cystic carcinoma of the head and neck: Predictors of morbidity and mortality. Archives of Otolaryngology–Head & Neck Surgery 125, 149–152 (1999).

46. van Weert, S., van der Waal, I., Witte, B.I., Leemans, C.R. & Bloemena, E. Histopathological grading of adenoid cystic carcinoma of the head and neck: analysis of currently used grading systems and proposal for a simplified grading scheme. Oral oncology 51, 71–76 (2015).

47. Wolbach, S.B. & Howe, P.R. Tissue changes following deprivation of fat-soluble A vitamin. The Journal of experimental medicine 42, 753–777 (1925).

48. Adriance, M.C., Inman, J.L., Petersen, O.W. & Bissell, M.J. Myoepithelial cells: good fences make good neighbors. Breast Cancer Research 7, 1–8 (2005).

49. Sirka, O.K., Shamir, E.R. & Ewald, A.J. Myoepithelial cells are a dynamic barrier to epithelial dissemination. The Journal of cell biology 217, 3368–3381 (2018).

50. Hanna, G.J., ONeill, A., Cutler, J.M., Flynn, M., Vijaykumar, T., Clark, J.R., Wirth, L.J., Lorch, J.H., Park, J.C. & Mito, J.K. A phase II trial of all-trans retinoic acid (ATRA) in advanced adenoid cystic carcinoma. Oral oncology 119, 105366 (2021).

51. Huynh, T.T., Sultan, M., Vidovic, D., Dean, C.A., Cruickshank, B.M., Lee, K., Loung, C.-Y., Holloway, R.W., Hoskin, D.W. & Waisman, D.M. Retinoic acid and arsenic trioxide induce lasting differentiation and demethylation of target genes in APL cells. Scientific reports 9, 1–13 (2019).

52. Jamieson, C.H., Ailles, L.E., Dylla, S.J., Muijtjens, M., Jones, C., Zehnder, J.L., Gotlib, J., Li, K., Manz, M.G. & Keating, A. Granulocyte–macrophage progenitors as candidate leukemic stem cells in blast-crisis CML. New England Journal of Medicine 351, 657–667 (2004).

53. Querfeld, C., Nagelli, L.V., Rosen, S.T., Kuzel, T.M. & Guitart, J. Bexarotene in the treatment of cutaneous T-cell lymphoma. Expert opinion on pharmacotherapy 7, 907–915 (2006).

54. Tallman, M.S., Andersen, J.W., Schiffer, C.A., Appelbaum, F.R., Feusner, J.H., Woods, W.G., Ogden, A., Weinstein, H., Shepherd, L. & Willman, C. All-trans retinoic acid in acute promyelocytic leukemia: long-term outcome and prognostic factor analysis from the North American Intergroup protocol: Presented in part at the 39th meeting of the American Society of Hematology, New Orleans, LA, December 1999. Blood, The Journal of the American Society of Hematology 100, 4298-4302 (2002).

55. Theodosiou, M., Laudet, V. & Schubert, M. From carrot to clinic: an overview of the retinoic acid signaling pathway. Cellular and molecular life sciences 67, 1423–1445 (2010).

56. Shalita, A.R. Mucocutaneous and systemic toxicity of retinoids: monitoring and management. Dermatology 175, 151–157 (1987).

57. Zhang, Y., Parmigiani, G. & Johnson, W.E. ComBat-seq: batch effect adjustment for RNA-seq count data. NAR Genomics and Bioinformatics 2 (2020).

58. Crowley, L., Cambuli, F., Aparicio, L., Shibata, M., Robinson, B.D., Xuan, S., Li, W., Hibshoosh, H., Loda, M. & Rabadan, R. A single-cell atlas of the mouse and human prostate reveals heterogeneity and conservation of epithelial progenitors. Elife 9, e59465 (2020).

59. Wolf, F.A., Angerer, P. & Theis, F.J. SCANPY: large-scale single-cell gene expression data analysis. Genome biology 19, 1–5 (2018).

60. Rousseeuw, P.J. Silhouettes: a graphical aid to the interpretation and validation of cluster analysis. Journal of computational and applied mathematics 20, 53–65 (1987).

61. Eilenberger, C., Kratz, S.R.A., Rothbauer, M., Ehmoser, E.-K., Ertl, P. & Küpcü, S. Optimized alamarBlue assay protocol for drug dose-response determination of 3D tumor spheroids. MethodsX 5, 781–787 (2018).

62. Hather, G., Liu, R., Bandi, S., Mettetal, J., Manfredi, M., Shyu, W.-C., Donelan, J. & Chakravarty, A. Growth Rate Analysis and Efficient Experimental Design for Tumor Xenograft Studies: Supplementary Issue: Array Platform Modeling and Analysis (A). Cancer Informatics 13, CIN. S13974 (2014).

